# Scalable and Synthesizable DNA Library Design via Reinforcement Learning

**DOI:** 10.64898/2026.07.07.736931

**Authors:** Scott Sussex, Ema Borevković, Frederieke Lohmann, Ningning Chen, Elena Lüthi, Sai T. Reddy, Andreas Krause

## Abstract

Designing DNA libraries is a key challenge from drug design to protein engineering and synthetic biology. Modern generative models offer opportunities to navigate the design space and propose specific sequences predicted to be effective in-silico. Designing *deterministic* libraries of specific sequences is however limited by the cost of DNA synthesis – the *synthesis barrier*. In contrast, high-throughput multiplexed screening can measure the function of billions of biological sequences in parallel. Harnessing this technology requires the design of randomized libraries with specific design constraints to achieve low synthesis costs. In practice, such stochastic libraries are often chosen heuristically, sacrificing control for scale. Is there a way to bridge AI-based in-silico sequence design with high-throughput experimentation? In this work, we introduce *Policy Gradients for Library Design (PGLD)*. PGLD uses a *synthesis-aware* parameterization of stochastic DNA libraries and optimizes them against a specified objective function. This allows for designing massive, controlled libraries without being limited by synthesis costs. We show how PGLD enables lab-in-the-loop design of multi-round high-throughput experiments, and large-scale in-vitro DNA sampling from generative models. Finally, we use PGLD to design a library of ∼ 10^6^ unique sequences at a cost of ∼ 700 USD to explore the mutation space of a broadly neutralizing influenza antibody.

## 1 Introduction

While increasingly powerful AI models can propose large numbers of biological sequences insilico, experimentally validating designs at scale remains a bottleneck. High-throughput multiplexed assays can screen potentially billions of sequences for functionality simultaneously, but explicitly synthesizing a library of this size is economically infeasible.

To reduce synthesis costs, randomized approaches like degenerate codon libraries are often employed. These approaches design a single DNA ‘template’, where mixes of nucleotides define a categorical distribution over nucleotides at each position (Fig. 1). The resulting libraries can generate huge diversity for the cost of synthesizing effectively a single DNA template. However, these methods offer limited control over the resulting library. To offer more control, Weinstein et al. [44] reframe degenerate synthesis as one instance of a broader family of stochastic synthesis methods, each of which takes a set of synthesis parameters to define a distribution over sequences which can then be optimized. Their proposed algorithm, Variational Synthesis, optimizes those parameters via maximum likelihood estimation on a pre-sampled target dataset.

**Figure 1:**
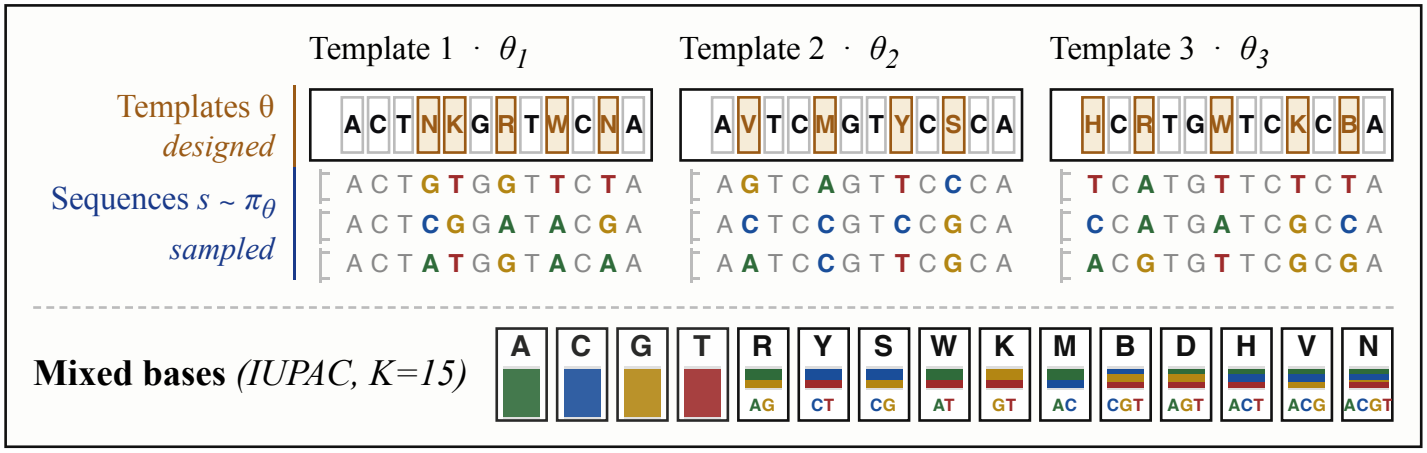
Stochastic DNA synthesis. PGLD optimizes a policy defined by *M* templates. A template is a sequence of chosen mixed DNA bases applied, in order, in its own reaction column. Examples of 3 possible synthesized DNA sequences from each template are shown below the template design. The sequences from each column are mixed together to form the final synthesized library. In our experiments, we show examples ranging from 1 to 1024 templates.

In this work, we approach library design as policy optimization of the synthesis distribution using methods from reinforcement learning. We introduce Policy Gradients for Library Design (PGLD), an algorithm which directly optimizes multi-template stochastic synthesis processes against a specified objective function. Similarly to Weinstein et al. [44], we parameterize all opportunities for randomization of a manufacturable synthesis process (see Fig. 1) and simulate it on a computer. PGLD iteratively samples libraries from the current parameters, scores them with an in-silico objective function, and updates parameters via policy gradients.

While previous work [56] has explored optimizing single template synthesis protocols via policy gradients, we introduce techniques for optimizing multi-template protocols. In several experiments across modalities and objectives, we demonstrate how scaling the number of templates can induce higher-performing libraries. A key advantage of our parameterization compared to prior work is that it results in multi-template libraries which can be affordably manufactured from commercial synthesis providers.

Furthermore, using direct on-policy optimization allows us to customize the objective function, and therefore adapt PGLD for various ML-guided design tasks. We demonstrate PGLD across diverse modalities (DNA aptamer, peptide binder, protein design) in a lab-in-the-loop setting to design sequential, multi-round high-throughput experiments. We also demonstrate how distribution matching can be cast as PGLD optimization to perform approximate in-vitro sampling from generative sequence models. Finally, we present an experimental case study using PGLD to design ∼10^6^ unique sequences for exploring the mutation space of a broadly neutralizing influenza antibody. The library is guided by a prediction model trained on local mutation data for binding two different strains. We validate experimentally via yeast surface display that the synthesized library, with higher mutational distances, is substantially more enriched for binding sequences than a representative heuristic library design approach.

## 2 The Stochastic Library Design Problem

We aim to design a library of *B* DNA sequences *S* = {*s*_0_, …, *s*_*B*−1_} each of length *L* over the alphabet *A* = {*A, C, G, T* }. We typically want to design the set *S* such that obtaining experimental labels for those sequences is maximally informative. In practice this will likely involve covering diverse, novel sequences that score highly under an in-silico fitness predictor *f* (*s*). Abstractly, the goal is to design *S* that maximizes an in-silico batch scoring function *R*(*S*). We provide some concrete examples in Section 4. For protein engineering applications, we can consider objective *R* that scores the corresponding protein sequences for DNA in *S*. Because deterministically synthesizing uniquely specified sequences is cost-prohibitive for large *B*, we instead optimize over the parameters of a stochastic synthesis process *π*_*θ*_:

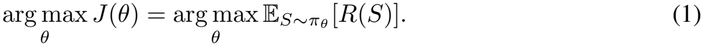

Here we parameterize *π*_*θ*_ as degenerate oligo pool synthesis [44], depicted in Fig. 1. The parameters *θ* consist of *M* template sequences, each of length *L*. A mixed nucleotide **v**^*k*^ ∈ Δ^3^ defines a categorical distribution over *A*. At position *l* ∈ [*L*] on a template *m* ∈ [*M* ], 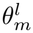 specifies an independent mixed nucleotide from a fixed set 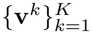 (in our experiments: the 15 standard IUPAC bases [7]).

To sample a sequence *s* ∼ *π*_*θ*_, we uniformly select a template *m* and independently sample a nucleotide at each position *l* based on 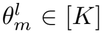. Let *S* ∼ *π*_*θ*_ denote sampling *B* such sequences *i*.*i*.*d*.. *θ* specifies a stochastic DNA synthesis protocol that can be ordered from commercial DNA synthesis providers. The model *π*_*θ*_ is an abstraction of the physical DNA synthesis process, which may in practice have, e.g., slightly different coupling rates for different pairs of nucleotides. Weinstein et al. [45] have previously shown experimentally that samples from an abstract synthesis policy *π*_*θ*_ closely align with the physical DNA samples synthesized by a protocol *θ*. Alternative stochastic synthesis methods are possible (cf. Appendix B) and also compatible with PGLD.

The cost of stochastic synthesis scales with the number of templates, each of which may produce many unique sequences, whereas deterministic synthesis produces one sequence per template. For this reason, deterministic synthesis rarely appears in the literature for designing libraries with over 10^6^ unique sequences.

## 3 Policy Gradients for Library Design

### 3.1 Optimizing Stochastic Synthesis Policies

Because the parameters 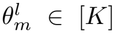 are discrete, we reparameterize *π* using continuous logits 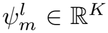 for each parameter, where 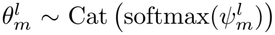. The *ψ* also define a probability distribution over sequences. Let *τ* denote a sampled batch containing both sequences 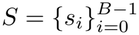 and the template 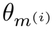 that produced each, where *m*^(*i*)^ denotes the index of the template that the *i*th sequence was sampled from. Since the sampling of DNA sequences is discrete, we use the score gradient estimator (REINFORCE [47]) to compute gradients: ∇_*ψ*_*J* (*ψ*) = E_*τ*∼*ψ*_[(*R*(*S*) − *b*)∇_*ψ*_ log ℙ [*τ* | *ψ*]] where *b* is a baseline. The log-likelihood of the batch admits the following form:

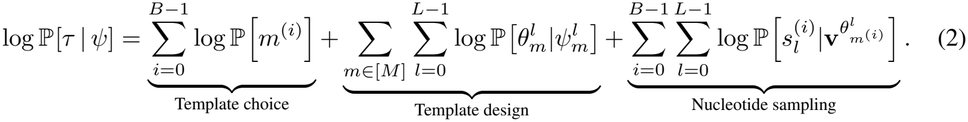

The template design term is the only one that conditions on *ψ*. We give the full algorithm in Algorithm 1 for the case where a single Monte-Carlo sample *N*_*MC*_ = 1 is used for each optimizer step. At convergence, we recover physically realizable parameters *θ* by taking the mode of each Cat 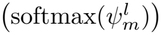. Theorem 1 (Appendix C) shows that the supremum of *J* over *ψ* equals the maximum over *θ*.

### 3.2 Scaling to Large Numbers of Templates

Algorithm 1 assigns a score to the entire batch and uses it to update all parameters. As the number of templates *M* increases, any individual template’s contribution towards the total reward becomes small, leading to high variance gradient updates. We reduce variance by assigning credit to each template individually. For many experiment design objectives (see Section 4.1), the marginal impact of each template on the global objective value diminishes as we increase *M*, making our objective a submodular set function over template designs [24]. Maximization of submodular set functions can be solved provably near-optimally by framing optimization as distributed welfare games [26], which motivates our practical use of specific per-template credit assignment strategies. We follow one of two strategies for decomposing the reward. Let components of *ψ* that relate to only template *m* be 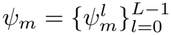. Let *S*_*m*_ denote sequences in a batch sampled by template *m* and *S*_\*m*_ = *S* \ *S*_*m*_.

#### Algorithm 1

Policy Gradients for Library Design (PGLD)

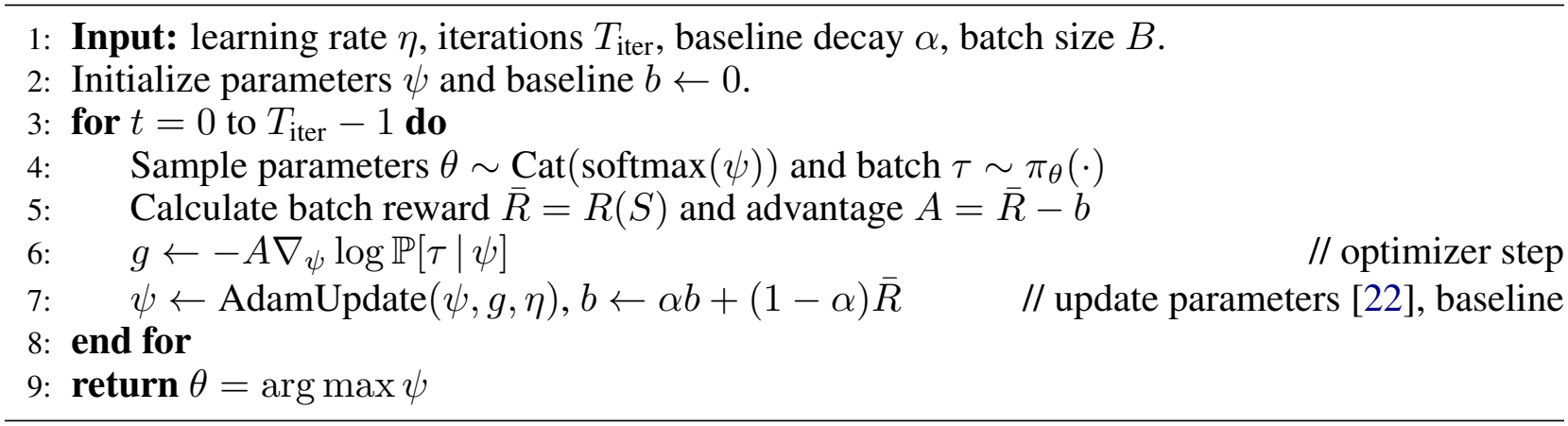

#### Equal Share

This can be applied to objective functions that sum up per-sequence rewards *r*(*s*): *R*(*S*) = Σ_*s*∈Unique(*S*)_ *r*(*s*). Template *m* optimizes the objective

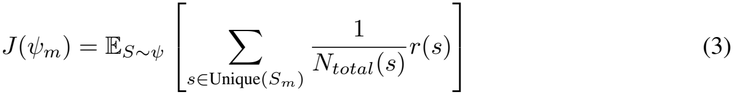

where *N*_*total*_(*s*) is the number of templates that sample *s* at least once. When equal share can be applied, it is straightforward to implement and efficient to compute.

#### Marginal Gain

This applies to more general rewards than equal share. It computes the difference between the global reward and the counterfactual reward if template *m* didn’t contribute:

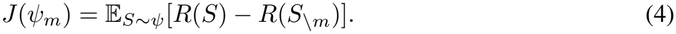

In both cases we effectively run *M* instances of PGLD, with each instance optimizing a single template with respect to its own objective. We give full pseudo-code in Appendix D.

## 4 Applications

So far we have described an algorithm that is largely agnostic to the chosen objective function. Here, we demonstrate how different objective functions enable AI-guided library design.

### 4.1 Steering Library Design with AI Models

When a surrogate model *f* (*s*) predicting sequence fitness is available, PGLD can optimize libraries to maximize diverse coverage of high-fitness sequences. We propose two primary objectives for achieving this:

**Mean-Entropy** arg 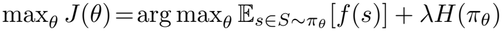, originally proposed in Zhu et al. [56], where *H* is the Shannon entropy and *λ* ∈ ℝ is a hyperparameter. This cannot be easily decomposed as a sum of per-sequence rewards, so we pair it with marginal gain for credit assignment. *H*(*π*_*θ*_) can be computed in closed form when *M* = 1, and for *M >* 1 must be approximated by a Monte Carlo estimate.

### Expected Hits

Given a specified fitness threshold *ρ*, a ‘hit’ is a sequence where *f* (*s*) *> ρ*. We optimize the expected count of unique hits

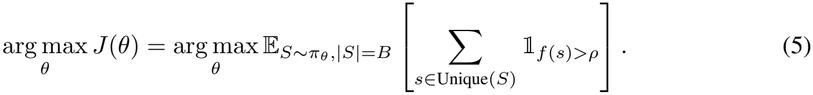

Expected Hits can be combined with the equal share rule when assigning credit per-template. For both objectives, when *B* is large and *f* is expensive to query, we can use importance sampling to give an unbiased estimate of the objective while only querying *f* with a subset of the batch at each iteration (as demonstrated in Appendix H). We study Mean-Entropy and Expected Hits as two possible objective functions that could be combined with PGLD, but emphasize that significant customization is possible. In the above objectives, for protein engineering tasks we compute uniqueness and entropy over DNA sequences with respect to their corresponding amino acid sequences.

### 4.2 Sequential Experiment Design

In multi-round lab-in-the-loop campaigns, at round *t* a surrogate model 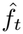 is trained on historical data H_*t*−1_ containing all sequences and fitness values observed in prior rounds [54, 27]. A typical goal is to discover a diverse range of high-fitness sequences. Achieving this requires formalizing objectives that can 1) use 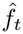 as a predictive model, and 2) discount sequences observed in previous rounds to encourage exploration of novel sequence space [41]. We can adapt the Expected Hits objective accordingly to optimize Cumulative Unique Hits:

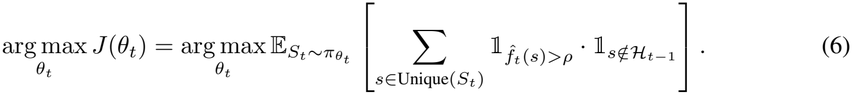

The full algorithm for designing lab-in-the-loop experiments with Eq. (6) is shown in Appendix E.

### 4.3 Sampling from a Generative Model In-Vitro

Sampling from an in-silico generative model [15, 8, 5] *p*_model_ (e.g., a protein language model) has been successfully used to propose experiments [16]; however, the explicit synthesis of millions of these individual samples for high-throughput applications imposes prohibitive costs. Each synthesized sample from *p*_model_ corresponds to one additional template, coupling the number of samples to the synthesis cost.

We can instead use PGLD to design a stochastic library that approximates *p*_model_ *in-vitro* by substituting log *p*_model_ for *f* (*s*) in the Mean-Entropy objective. This objective corresponds to minimizing the reverse KL divergence from the tempered distribution 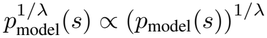, that is

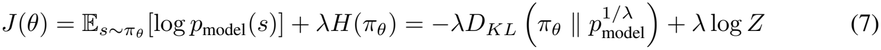

where *Z* denotes the normalizer of 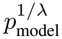 which does not depend on *θ*.

Furthermore, we can enforce constraints on the library C (e.g., a maximum edit distance from a starting sequence, no early stop codons), by assigning a negative penalty *η* to constraint-violating sequences:

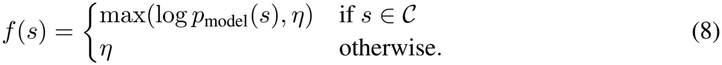

## 5 Experiments

### 5.1 PGLD as an Optimizer

We first assess the ability of PGLD to optimize libraries on seven different fitness landscapes. Five are based on smaller domains exhaustively enumerated via high-throughput screening, and two are based on strong machine learning models.

Since some landscapes have small domains (most 8 to 12 nucleotides), we validate with small batch sizes (order of *B* = 10^3^); however, in our applications in Section 5.3 and Section 5.4 we demonstrate PGLD on batch sizes of 10^6^ or more. We show results on a subset of domains here, with experiments on all domains shown in Appendix G.

Using the Expected Hits objective, we find that PGLD consistently optimizes for libraries that contain many more high-fitness sequences than random NNK/NNN libraries (Fig. 2a). While restricting the chemistry to standard IUPAC bases instead of arbitrary ratios of nucleotides (see Appendix B) reduces performance for a single template library, it is by far the most practical for scaling to larger numbers of templates. Using just a few (eight) templates is sufficient to achieve most of the attainable performance across all landscapes, with continued improvements as template count is increased further (Fig. 2b). In Fig. 2c we see that the per-template assignment of credit using equal share accelerates optimization compared to a global optimizer. The Mean-Entropy objective is demonstrated in our experimental work in Section 5.4. Unless stated otherwise, experiments in subsequent sections use PGLD with only IUPAC bases.

**Figure 2:**
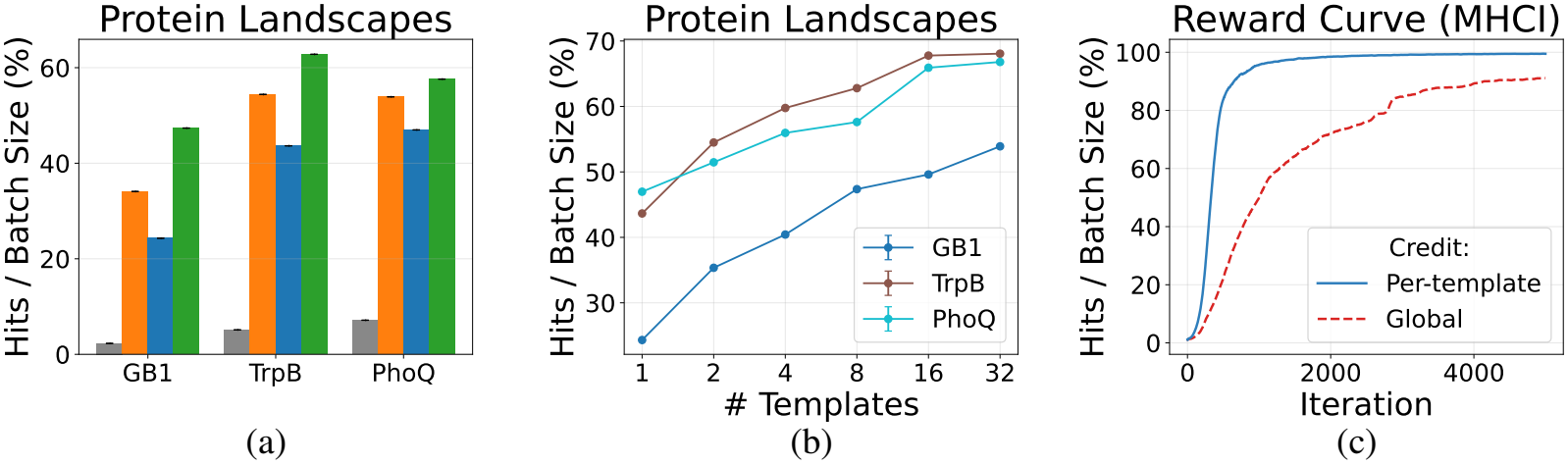
The PGLD optimizer, expected hits objective. **a.** PGLD produces libraries with high objective value on 3 protein landscapes, particularly when increasing the number of templates used. We compare (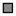) Random NNK; (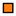) PGLD (Arbitrary Base, 1 Template); (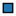) PGLD (IUPAC, 1 Template); (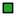) PGLD (IUPAC, 8 Templates). **b**. On the same fitness landscapes, scaling the template count for PGLD with IUPAC leads to improved solutions. **c**. On the MHCI landscape with 32 templates, template-level credit assignment via equal share leads to substantially more efficient optimization than global credit assignment.

### 5.2 In-Silico Lab-in-the-Loop Experiments

To evaluate PGLD for multi-round experiment design, we simulate a 5-round sequential campaign across our seven empirical fitness landscapes, all starting with a random NNK/NNN round. While in Section 5.1 we directly optimize on the empirical fitness landscapes, here we assume the landscape is completely unknown a priori and that we must perform sequential experiments to discover high fitness sequences. Our simulated campaigns assume noiseless fitness measurements (we query the empirical fitness landscape to measure ground truth), no batch effects, and smaller batch sizes (10^3^ − 10^4^) to account for the small domain sizes of the empirical fitness landscapes.

As a surrogate model for PGLD, at each round we train a convolutional neural network (CNN) 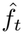 on all historically observed sequence-fitness pairs. We optimize the Cumulative Unique Hits objective in Eq. (6) with a 50-template design. We compare against two baselines: 1) continued random sampling, and 2) a rational design approach [28, 42], which is representative of strategies that combine the most promising single mutations combinatorially to form a library [46, 14]. As in Mason et al. [28], for protein engineering tasks we compute target amino acid ratios at each position based on how often each has appeared in hits, and then select a single IUPAC template that best approximates these target distributions. For DNA sequence design tasks an analogous strategy is implemented.

Across all landscapes, PGLD significantly accelerates the discovery of novel hits (Fig. 3). On representative landscapes such as TrpB, a 4-amino-acid protein landscape, and Switch, a 10-nucleotide DNA landscape, PGLD yields over a 100% improvement in cumulative hits compared to baselines by round two. Over all 5 rounds, PGLD achieves at least a 30% improvement in cumulative hits across all landscapes. Experiments are repeated over 10 random seeds and averaged, with standard errors shown.

**Figure 3:**
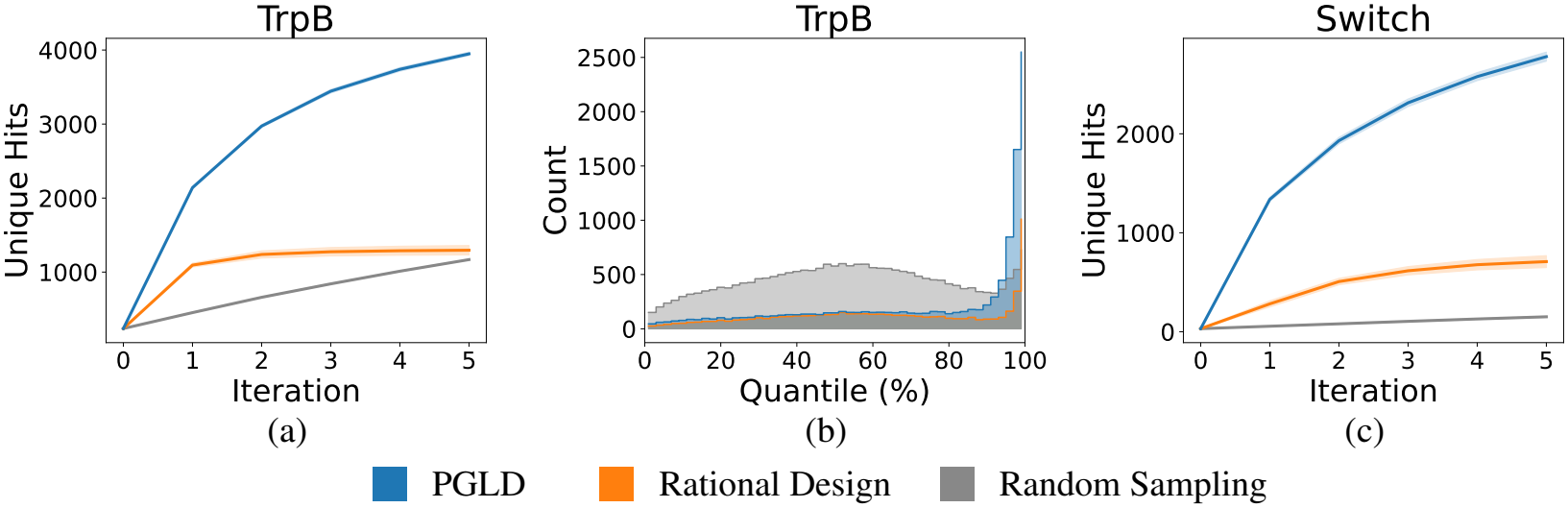
In-silico lab-in-the-loop experiments. For each of the seven empirical fitness landscapes we provide comparisons of PGLD, rational design and random sampling (NNN/NNK). **a**. The cumulative hits across rounds for the TrpB protein landscape. **b**. The quantile distribution of all unique, valid sequences sampled. All fitness measurements in the ground-truth datasets are mapped to quantiles for visualization. **c**. The cumulative hits across rounds for the Switch DNA landscape.

Analyzing the distribution of sampled sequences (e.g., Fig. 3b) reveals that PGLD with the Expected Hits objective concentrates library density in the upper tail of the fitness distribution, and not just on sequences barely above the set threshold for a hit. Details on the training of 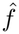, baseline implementations, and results for all landscapes are provided in Appendix I.

### 5.3 Library Design for Sampling from Generative Models In-Vitro

We apply PGLD to design 1024-template libraries that approximate sampling from protein language models (PLMs) in-vitro for two antibody CDRH3 design tasks: Trastuzumab (using the autoregressive IgLM [38]) and the broadly neutralizing influenza antibody C05 (using the masked ESM2-150M [25]). Trastuzumab has an 11-amino-acid CDRH3, and C05 has a 24-amino-acid CDRH3. For C05, we enforce a maximum edit distance of 6 from the wild-type (WT) sequence using Eq. (8), and in both experiments penalize early stopping. Since the reverse KL is highly mode-seeking [3], we set *λ >>* 1 to prevent mode collapse.

We compare with Variational Synthesis [44, 45], the state-of-the-art method for learning stochastic DNA synthesis policies that match generative models. Variational Synthesis fits synthesis parameters to a pre-sampled dataset (we use approximately 8 × 10^6^ samples for IgLM and 4 × 10^6^ for ESM2) via Expectation-Maximization to minimize the forward KL divergence, as opposed to the tempered reverse KL divergence we optimize with PGLD in Eq. (7). IgLM can be sampled from autoregressively (we retain only samples of length 11), and for ESM2 we use Gibbs sampling. Sequence likelihoods can be computed exactly for IgLM and estimated with pseudolikelihoods [36] for ESM2. Both Variational Synthesis and PGLD use a degenerate oligonucleotide synthesis with 1024 templates and 15 mixed bases. Weinstein et al. [45] show that there is only a small discrepancy in distribution between in-vitro sampled sequences and those sampled from degenerate oligonucleotide libraries in-silico; therefore, we evaluate the two methods in-silico.

To match the original implementation released by Weinstein et al. [44], we allow Variational Synthesis to learn both the nucleotide ratios of each mixed base (Arbitrary Base) and the relative weighting of each template (uniform for PGLD). Compared to PGLD, this allows Variational Synthesis the advantage of optimizing over a more expressive class of synthesis policies. To the best of our knowledge, this parameterization is not realizable with an off-the-shelf product from a commercial DNA synthesis provider. Modifying the Variational Synthesis parameterization to use IUPAC and uniform template weights leads to significantly degraded performance.

After optimizing a PGLD library and Variational Synthesis model, we sample sequences and compare them to unseen sequences from the PLM (Fig. 4). The PGLD-optimized Trastuzumab library has a theoretical diversity (the number of unique sequences that could be sampled from the library) of 1.5 × 10^13^, and the C05 library has a theoretical diversity of 3 × 10^9^. In both settings, the PGLD libraries contain mostly sequences with a high likelihood under the PLM, broadly cover the PLM distribution in UMAP [29] space, and can strictly follow constraints. In Table 1 we compute various quantitative metrics for how well PGLD and Variational Synthesis libraries match the reference distribution [45]. PGLD more reliably produces libraries that obey hard constraints. For distribution-matching metrics, the relative ranking of PGLD and Variational Synthesis depends on the setting and metric considered.

**Table 1:** In-vitro sampling from generative models. We compare 10^5^ sequences from PGLD and Variational Synthesis libraries against the target PLMs. Distributional metrics are described in Section J.4.

| **Library** | **MMD<sup>2</sup> ↓** | **Witness (%) ↑** | **BEAR BF ↓** | **Eff. Diversity** |
|---|---|---|---|---|
| *Trastuzumab (IgLM)* |  |  |  |  |
| PGLD | 0.00122 | 34.3 | 1.36 | 7.5 × 10<sup>8</sup> |
| Var-Synth (all / valid only) | 0.00299/0.00048 | 37.2/41.1 | 1.11/1.10 | 1.5 × 10<sup>12</sup>/– |
| *C05 (ESM2)* |  |  |  |  |
| PGLD | 0.00093 | 28.2 | 0.512 | 1.5 × 10<sup>9</sup> |
| Var-Synth (all / valid only) | 0.00971/0.00147 | 29.0/22.7 | 0.623/0.499 | 8.0 × 10<sup>12</sup>/3.0 × 10<sup>10</sup> |

**Figure 4:**
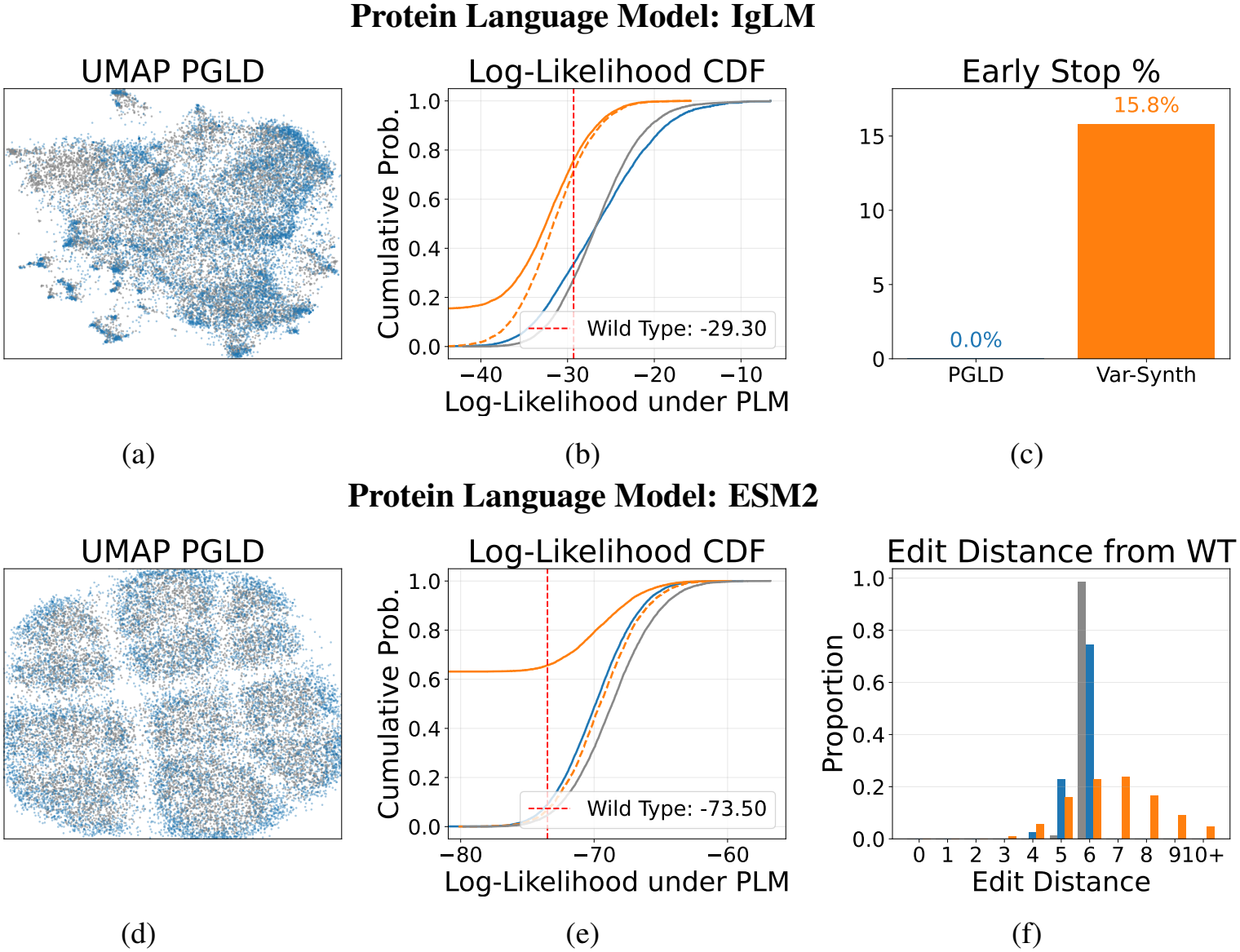
In-vitro sampling from generative models. We compare sequences sampled from PGLD libraries (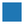) and Variational Synthesis models (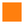, dashed = valid-only) against the target PLMs (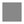) . **(Top Row) Target: IgLM. a**. For Trastuzumab CDRH3s, PGLD covers the target distribution in UMAP space and **b**. concentrates density on high-likelihood sequences. **c**. PGLD avoids early stopping codons. **(Bottom Row) Target: ESM2. d**. For C05 CDRH3s under a maximum edit distance of 6 from the wild type, PGLD covers the target distribution in UMAP space and **e**. concentrates density on high-likelihood sequences. **f**. PGLD obeys the edit distance constraint.

The qualitatively different behavior stems from the two different objectives: the forward KL objective of Variational Synthesis attempts to cover the full support of its training samples, but will do so despite possibly high rates of invalid, early stopping, or low-likelihood sequences. Using a reduced penalty for invalid sequences in PGLD might allow one to trade off rates of sequence validity against improved metrics on a valid-only subset.

Designed Trastuzumab templates under PGLD have significant variations in theoretical diversity (Fig. 12). To use them in a downstream application, e.g., a display screen, it would be desirable to produce more balanced template diversities and prevent oversampling of certain sequences. Since here the singular goal is to match a PLM distribution, the imbalanced template behavior is favorable because it allows the PGLD-learned libraries to place high-likelihood sequences on less diverse templates, thereby increasing the probability mass assigned to them.

### 5.4 Lab Validation: Influenza Antibody Library Design

As an experimental case study, we apply PGLD to design a CDRH3 library for the broadly neutralizing influenza antibody C05 [11], and screen it with yeast display. Previous antibody engineering efforts have observed a trade-off between affinity and breadth of binding [49], but only mutated a small subset of the 24 CDRH3 residues. We use PGLD to generate a library of C05 mutants considering mutations to any CDRH3 residue, aiming to retain binding to two influenza strains of subtypes H3N2 and H1N1. Such a library can be screened to map out sequence-activity relationships for C05 mutants.

We first obtain a double deep mutational scan (DDMS) [12] dataset measuring log-enrichment values (a proxy for binding) for all single and double mutants against the two influenza strains using yeast display [4]. We fine-tune ESM2 to predict log-enrichment from sequence (see Appendix K) to obtain separate models *f*_*H*1*N*1_, *f*_*H*3*N*2_.

To design libraries we optimize the Mean-Entropy objective, using *f* = *f*_*H*1*N*1_ + *f*_*H*3*N*2_. We sweep *λ* for PGLD with different template counts and the single-template, arbitrary-base algorithm of Zhu et al. [56]. We choose a target theoretical diversity of 10^6^ based on the expected throughput of our yeast display assay. In Fig. 5a we circle the library used experimentally, a 64-template library with a theoretical diversity of ∼ 9 × 10^5^. This library has a modal edit distance of 6 from the wild type (WT) and a maximum of 11 (Fig. 5b). The library’s distribution over amino acid sequences has an effective diversity (exponential of entropy) of 2.39 × 10^5^.

**Figure 5:**
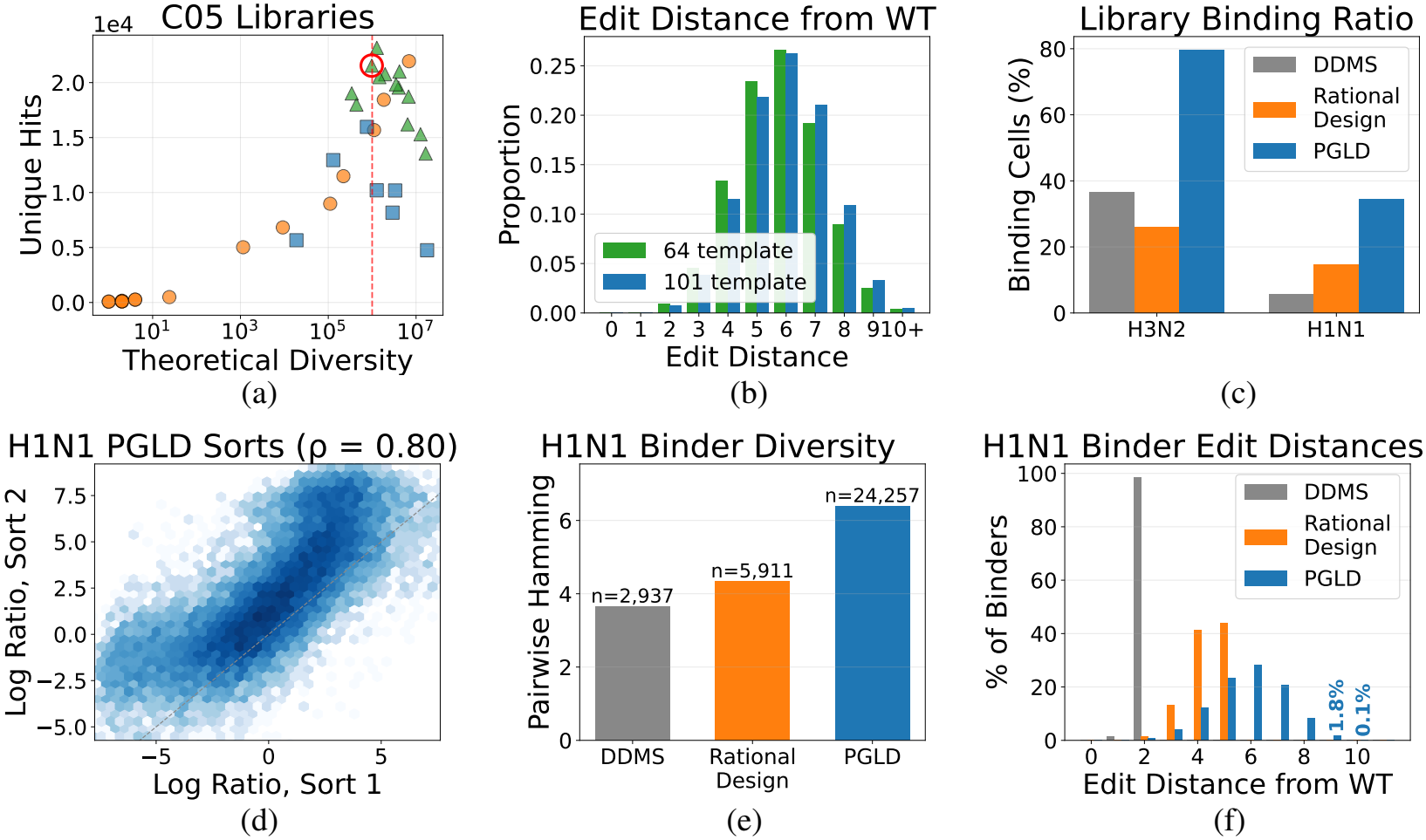
C05 antibody engineering. **a.** PGLD libraries optimized with different # templates and *λ* values. Each point is a library, showing expected unique predicted hits from 10^5^ sequence samples (predicted log enrichment ≥ wild-type (WT) ground truth on both strains under the surrogate model). (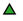) IUPAC, 64 templates; (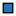) IUPAC, 8 Templates; (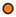) Arbitrary Base, 1 template (Zhu et al. [56]). The red dashed line represents the target diversity of the library, and the circled point is our final design. **b**. The in silico edit distance from wild type for samples in our final design with and without template duplication. **c**. The % binding population of yeast cells expressing the C05 mutant for a double deep mutational scan, a rationally designed library, and the PGLD library. **d**. Spearman’s correlation in log ratio scores for sequences with ≥ 40 pre-sort reads on both H1N1 PGLD library sorts. **e**. Binders to H1N1 in the PGLD library are diverse, measured by mean pairwise Hamming distance of binding sequences. **f**. PGLD H1N1 binders have large edit distances from the wild type.

We synthesize the PGLD library with a commercial synthesis provider for ∼ $700 and screen it with yeast display under the same conditions as the DDMS library. Before synthesis, we duplicate the templates with the highest theoretical diversity to cover the most rarely sampled sequences more evenly (see Appendix L), giving a 101-template library. When scored on the same in-silico metrics as Fig. 5a, the 101-template library has an estimated 2.1 × 10^4^ unique hits from 10^5^ samples (compared to 2.2 × 10^4^ for the 64-template library), a greater effective diversity of 4.00 × 10^5^ and a comparable edit-distance distribution shifted slightly towards higher edit distance (see Fig. 5b). We also screen a rationally designed library [28] that mutates up to 5 residues and is based on the DDMS data.

In Fig. 5c we plot the % binding population among yeast cells for each library, showing a 7× increase compared to the DDMS library on the more challenging H1N1 strain, and over a 2× increase compared to the rationally designed library. While this bulk-level readout does not resolve specific sequences’ affinities, it demonstrates that the PGLD library is substantially enriched for binders to both strains.

The workflow to obtain sequence-level binder data for each target is described in Appendix L. For H1N1, PGLD library binders show a high diversity (Fig. 5e). Many have a high mutational distance from the wild type (Fig. 5f), with several binders found that have a mutational distance of 10 (nearly half the amino acids in the CDRH3). Since the PGLD library has few H3N2 non-binders, there is only weak selection during sorting and the sequence-level data for this antigen is less reliable. This is discussed in Appendix L.

## 6 Related Work

Degenerate codon designs have long been used to overcome scaling limits in DNA synthesis. In practice most high-throughput libraries use undirected randomization (e.g., NNK) or rational design principles such as those we compare to in Section 5.2 [28]. PGLD differs from prior work in supporting both arbitrary objectives and expressive multi-template designs.

LibDesign [30], SwiftLib [19], and DeCoDe [37] design libraries that optimize coverage over a set of desired sequences. Zhu et al. [56] and Yang et al. [52] optimize library-level fitness-diversity objectives over single templates. Yang et al. [53] and Wu et al. [51] use machine learning to inform library designs where the target distribution of amino acids is specified independently at each position (as in the rational design benchmark we use). POCoM [43] maps a Pareto frontier over two competing objectives, using a single template. Weinstein et al. [44] also design DNA libraries by parameterizing multi-template DNA synthesis on a computer. They developed Variational Synthesis, which optimizes libraries to distribution-match a provided target dataset. An orthogonal approach to stochastic synthesis is deterministically synthesizing lists of chosen sequences, which has resulted in libraries of up to one million sequences [1, 10, 40], but cannot scale substantially further due to the cost of synthesis.

For lab-in-the-loop experiments, machine learning approaches such as batch Bayesian optimization [9] have been successfully applied [35]. These methods propose surrogate objective functions that trade off exploration and exploitation. PGLD is complementary, optimizing over a physically realizable synthesis policy instead of an unconstrained batch of sequences. Surrogate objective functions from Bayesian optimization could inform new objective functions in PGLD.

## 7 Discussion

PGLD addresses the longstanding trade-off between scale and control in DNA library design, unlocking new capabilities for using in-silico models to design massive experiments. We evaluate PGLD on in-silico benchmarks across diverse modalities, including aptamer, peptide, antibody, and protein design tasks. By directly optimizing stochastic synthesis policies against a custom objective function, PGLD surpasses relevant baselines on tasks ranging from sequential experiment design to in-vitro sampling from generative models. We demonstrate PGLD experimentally by synthesizing a diverse CDRH3 library for a broadly neutralizing influenza antibody and validating sequence fitness via yeast surface display. The designed library explores higher mutational distances than a heuristic baseline while discovering substantially more high-fitness sequences.

Many interesting directions remain. Commercial DNA synthesis providers cap sequence lengths at a few hundred nucleotides, so designing libraries with longer variable regions will require modeling the downstream assembly of subsequences. Co-designing synthesis hardware alongside library design algorithms could yield more controllable or economical stochastic synthesis protocols. Finally, many of our experiments focus on binding assays, where in-silico models are making rapid progress. Functional measurements beyond binding represent a substantially less explored frontier that would benefit from scalable, active experiment design.

## Acknowledgments and Disclosure of Funding

This research was supported by the Swiss National Science Foundation under NCCR Automation, grant agreement 51NF40 180545. This publication was created as part of NCCR Catalysis (grant number 225147), a National Centre of Competence in Research funded by the Swiss National Science Foundation.

Thank you to the Eric and Wendy Schmidt Center for hosting S.S and A.K during their visit there. Thank you to Caroline Uhler, Aldo Pacchiano and Jiaqi Zhang for relevant discussions while S.S and A.K were guests at the Eric and Wendy Schmidt Center. Thank you to Lars Lorch, Mojmir Mutny, Bhavya Sukhija and Fabrice Schlatter for relevant discussions with S.S at ETH. Thank you to Bhavya Sukhija, Miles Wang-Henderson, Lars Lorch, Louis Claeys, Teodora Kovacevic and Yi-Yi Ly for feedback on drafts of this work.

## A Code

Code for PGLD, the benchmark landscapes, baseline implementations and instructions to reproduce the in-silico experiments of section Section 5.1 to Section 5.3 are available at https://github.com/lasgroup/pgld.

## B Other Stochastic Synthesis Parameterizations

While we focus on degenerate oligo pools, other stochastic synthesis methods are possible [44] and many are compatible with PGLD.

### Arbitrary Base Synthesis

In arbitrary base synthesis, we are not restricted to a predetermined set of *K* mixed bases. For every position, the experimentalist can specify a different mixed base. In our notation this corresponds to an extreme case where the number of mixed bases is *K* = *M* × *L*. Algorithm 1 can be repurposed for this setting. Since this synthesis method relies on designing custom mixed bases for every template position, it is challenging to scale to large *M* in practice.

### Arbitrary Codon Synthesis

For protein design tasks, one can also design libraries on the codon level. In practice this is done by performing DNA synthesis with blocks of trinucleotides (nucleotide triplets). In our notation, this corresponds to defining a new alphabet *A*_codon_ = *A*^3^ and considering a sequence length *L*_codon_ = *L/*3. At each position on each template, an arbitrary distribution over the 64 trinucleotides can be specified. Algorithm 1 can be repurposed for this setting. Arbitrary codon synthesis has some benefits, such as guaranteed prevention of early stop codons. However it is even more challenging than arbitrary base synthesis to scale beyond a single template, since it relies on custom mixes of 64 possible trinucleotides at each position.

When simulating both of these arbitrary synthesis techniques on a computer, it will appear possible to select potentially extremely small probabilities of a nucleotide or trinucleotide appearing at a given position. In practice this is not possible to do accurately, since these probabilities correspond to physical concentrations of a nucleotide or trinucleotide and there will be some experimental error.

It is also possible to jointly learn the categorical distribution assigned to each **v** in degenerate oligonucleotide synthesis instead of pre-specifying them [44]. We do not study in this work how to achieve that with PGLD. In practice, we found that as of the time of writing, specifying the available mixed-bases to be all IUPAC bases was most compatible with what commercial DNA synthesis providers can offer for *M >* 1.

Other possible stochastic synthesis protocols are discussed in Weinstein et al. [44].

## C PGLD Optimizes for a Physically Realizable Synthesis Protocol

For a degenerate oligo pool, the logits *ψ* optimized over in PGLD define a distribution over mixed bases at each position in each template. This makes the mixed base at each position for a given template stochastic. A trained policy that outputs a stochastic choice over mixed bases at each position would be undesirable, since in practice we must submit a single mixed base for each position in each template (we must submit a design *θ* ∈ Θ). However, we can show that the supremum of the expected reward *J* (*ψ*) over the logit parameter space equals the maximum expected reward achievable by a deterministic design *θ* ∈ Θ.

### Theorem 1

(Optimal Mixed Base Choice). *The supremum of the expected reward over the logit parameter space equals the maximum expected reward achievable by a deterministic design:*

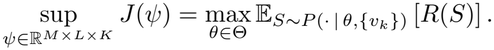

*Proof*. Assume that the available mixed bases 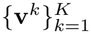 are fixed. The supremum of the expected reward is given by:

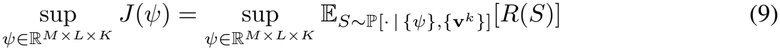

We can expand the expectation over sequences *S* by first taking the expectation over *θ*:

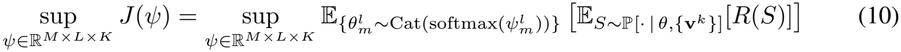

Because the expectation of a function over a random variable is always upper-bounded by the maximum value of that function (E_*X*_ [*f* (*X*)] ≤ max_*X*_ *f* (*X*)), we have:

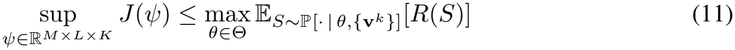

We have now demonstrated that the supremum of *J* (*ψ*) over logits *ψ* is upper bounded by the maximum expected reward over a deterministic design *θ*. What remains is to show that it is also lower bounded. Let 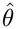 be the deterministic design achieving the maximum expected reward. For each *n* let’s define the parameters *ψ*^(*n*)^ such that the vector 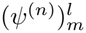 equals *n* for the mixed base choice 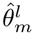 and zero otherwise. Since *θ* ∼ Cat(softmax(*ψ*)) we have:

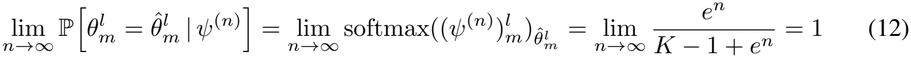

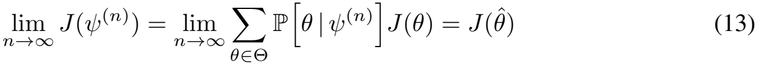

Eq. (13) follows because Θ is finite and the mixed base choices are independent. It therefore follows that:

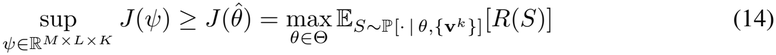

Having established both inequalities, we can conclude:

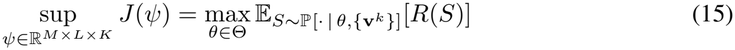

This completes the proof.

In practical cases where PGLD could converge to a *ψ* that corresponds to a stochastic allocation, in this paper we take the modal 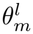 given 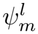 for all *l, m*. This doesn’t guarantee that *J* (*θ*) ≥ *J* (*ψ*), but in practice seems to work sufficiently well. If one insisted on ensuring that the final *θ* was chosen such that *J* (*θ*) ≥ *J* (*ψ*), one could design a sampling method that ensures this (greedy conditional sampling). Alternatively, one could repeat the sampling of *θ* and choose the one with the highest *J* (*θ*) to increase the likelihood of *J* (*θ*) ≥ *J* (*ψ*).

## D Further Details for Scaling Template Count

Here we provide full pseudocode for the methods described in Section 3.2 in Algorithm 2. For readability, we use *N*_MC_ = 1 but incorporating a larger number of MC samples per gradient update is straightforward. In the implementation of updating each template separately, we maintain a separate baseline for each template. When the loss has multiple components as in the Mean-Entropy objective, we in practice maintain a separate baseline for each of the two terms, for each template.

## E Sequential Experimental Design

Here we provide the full pseudocode for the method described in Section 4.2, for designing sequential experiments with PGLD.

### Algorithm 2

Multi-Template PGLD via Distributed Welfare Games

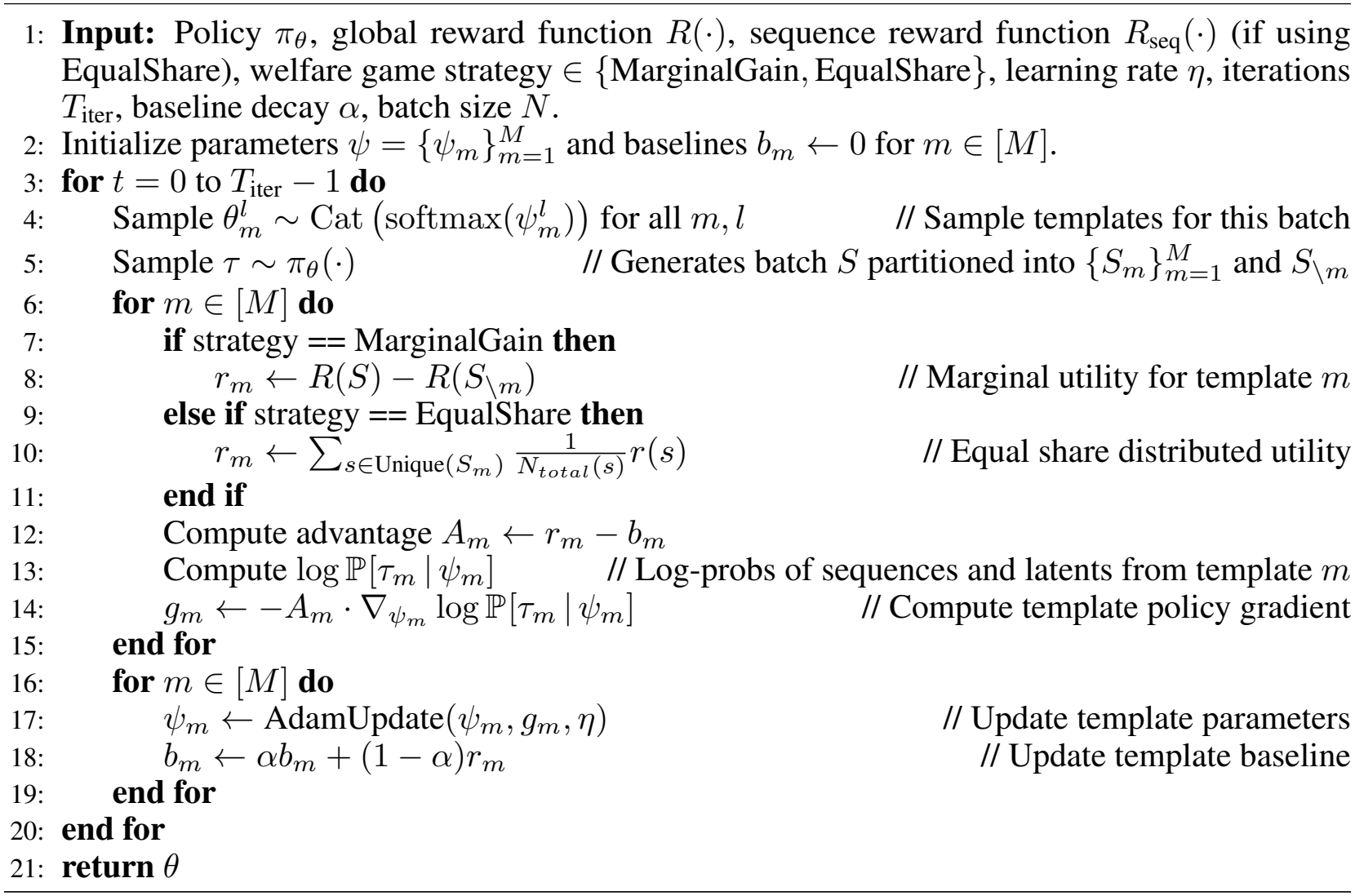

### Algorithm 3

Sequential Experiment Design

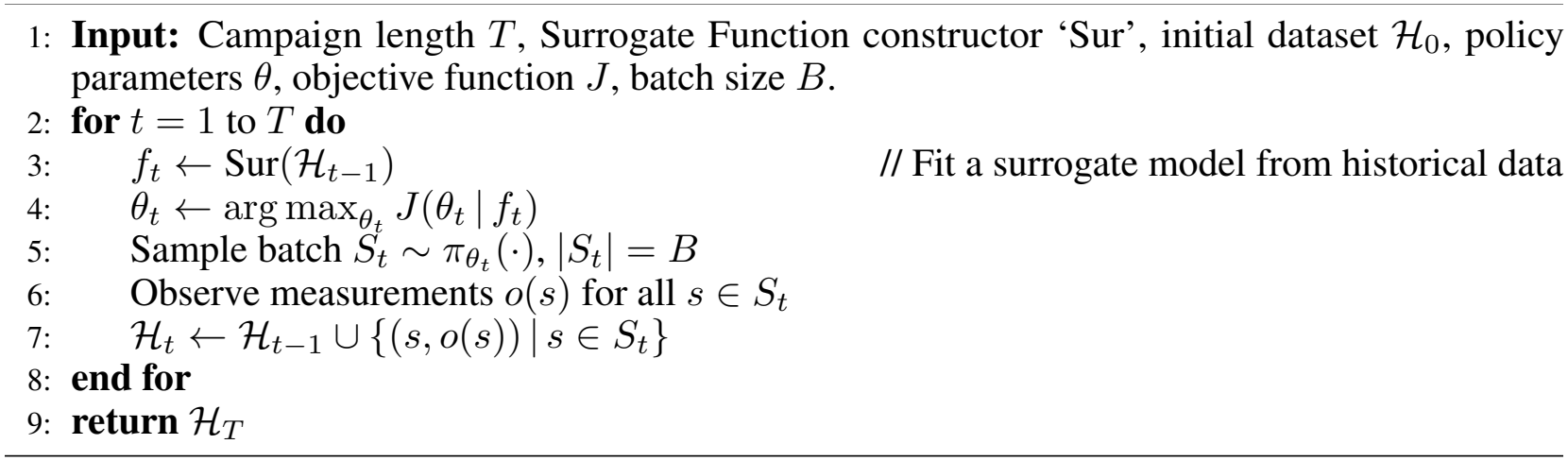

## F Experimental Settings

We source empirical fitness landscapes from two places. First, we consider two landscapes based on strong ML models trained on sequence data where the domain is densely sampled, but not all combinations exhaustively. Second, we consider landscapes with experimental measurements of DNA or protein sequences where for some subset of positions, all combinations of nucleotides or amino acids have been exhaustively scored.

### MHC class I binding affinity (MHCI)

O’Donnell et al. [31] train MHCflurry (Apache 2.0 license), a model predicting peptide-MHC class I binding affinity. We treat this model as a ground truth for simulating a high-throughput peptide design campaign where the goal is to discover diverse, high-affinity binders for a specific allele. We set the target to be the HLA-A*02:01 allele. This mimics real world experiments where yeast display has been used to screen combinatorial libraries of peptides for MHC affinity [17]. Our implementation of this landscape scores peptides of length 9 amino acids using MHCflurry. Throughout Section 5.1 we define the fitness of a peptide as 1 − *q/*100, where *q* is the MHCflurry predicted affinity percentile. Fitness therefore lies in [0, 1] and higher values correspond to higher predicted binding affinity.

### Molecular switch (Switch)

The task is to identify short DNA sequences, Switch Domains (SD), that can convert a pre-designed aptamer into a multi-component switch that undergoes a conformational change in response to a specific target small molecule. This has applications in biosensor design where the aptamer is hybridized to a fluorophore. The presence of the target molecule should trigger a structural change that ultimately results in a detectable change in fluorescence. Yoshikawa et al. [55] previously published data (CC BY 4.0) from screening a large number of random 10-bp DNA Switch Domains for ability to create a switch with a known aptamer for ATP. The data pairs each sequence with a relative fluorescence change measurement. We train a convolutional neural network model with a feed-forward head on this data and stop the training once the maximum Pearson’s *r* of 0.75 on a held-out validation set is reached. Training takes 7 hours on one Nvidia Quadro RTX 6000. Additional training hyperparameters are reported in Table 2. We then treat this model as ground truth in simulating a new screen where we aim to find 10bp DNA sequences that maximize the change in fluorescence when used in this molecular switch for ATP.

**Table 2:** Switch landscape log-fold change oracle training hyperparameters.

| **Hyperparameter** | **Value** |
|---|---|
| # Convolution layers | 8 |
| # Feed-forward layers | 2 |
| Kernel size | 3 |
| Dropout probability | 0.1 |
| Optimizer | Adam |
| Learning rate | 10<sup>−4</sup> |
| Loss function | MSE |
| Batch size | 512 |
| Max epochs | 1000 |
| Validation split | 10% |

**Table 3:**
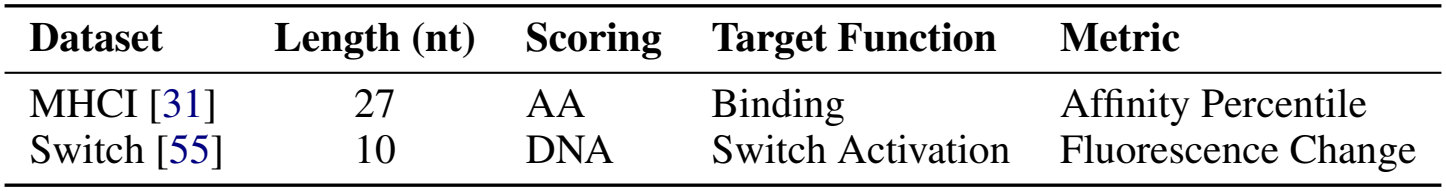
Landscapes based on querying an ML model.

**Table 4:**
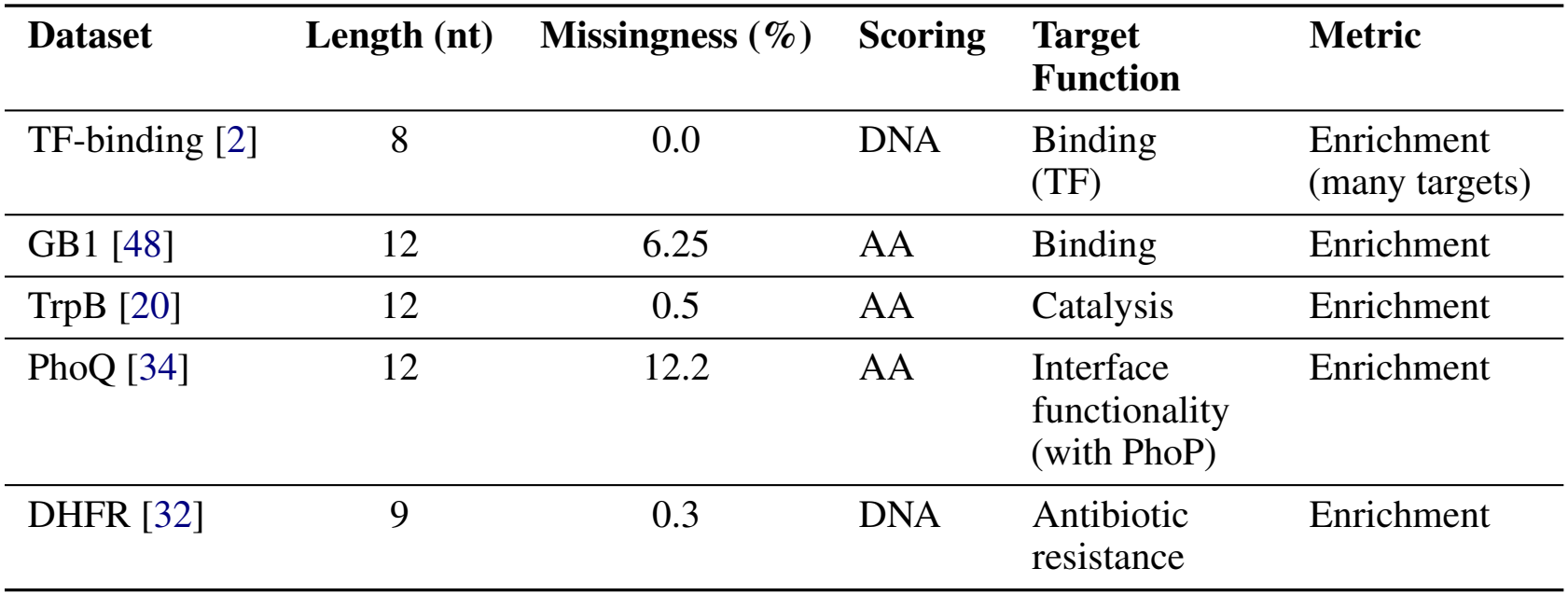
Landscapes based on querying real data.

We create several landscapes on datasets where the fitness landscape for some subset of nucleotide or amino acid positions is near exhaustively searched. In all cases, we use the dataset as ground truth to simulate a campaign exploring mutations which can improve fitness (however that is defined for that dataset). In all cases where we have exhaustive data on mutations at a few sites in a larger sequence, we design libraries that also only vary these positions. Any missingness is imputed by either assuming that sequence would score the minimum recorded value for the assay, or copying the imputation strategy used in the original work.

### TF-binding

Barrera et al. [2] screened all possible DNA sequences of length 8 for binding to 159 different transcription factors (TF) (license not found). This allows us to establish ground truth for a hypothetical screen of finding DNA aptamers that bind strongly to one of these transcription factors. The raw datasets were extracted from the FlexS package [39] (Apache 2.0).

### GB1

Wu et al. [48] study the fitness landscape of protein GB1. For all possible amino acid combinations and 4 relevant sites, they estimate binding affinity to IgG-Fc using mRNA display. This leads to a dataset (CC BY 4.0) of 160k different sequences.

### TrpB

(Creative Commons Zero) Johnston et al. [20] map the fitness landscape of an enzyme TrpB in a nonnative environment, varying 4 amino acid residues in the active site exhaustively (160k total combinations). Fitness is quantified as ability to catalyze the native reaction for TrpB.

### PhoQ

Podgornaia and Laub [34] exhaustively screened 4 amino acid positions in the Escherichia coli protein kinase PhoQ that drive recognition of PhoP. They measured the ability for the mutant PhoQ to form a functional interface with PhoP (license not found).

### DHFR

Papkou et al. [32] exhaustively vary all combinations of 9 nucleotides (>270k combinations) of the Escherichia coli metabolic gene folA (CC BY 4.0). The varied positions can confer resistance to an antibiotic. Fitness was defined and measured based on antibiotic resistance.

The tables below provide a summary of the experimental settings we use to study PGLD. For ‘length’, we report the number of DNA bases that can be varied, though there may be bases in-between these that are fixed. Scoring determines whether the data or model evaluates the DNA sequences designed or the amino acid sequence they are transcribed and translated to.

## G PGLD as an Optimizer

Here we describe the full experimental setup and results for Section 5.1. In Fig. 6 we show the results for the Expected Hits objective from Section 5.1 but for all landscapes.

**Figure 6:**
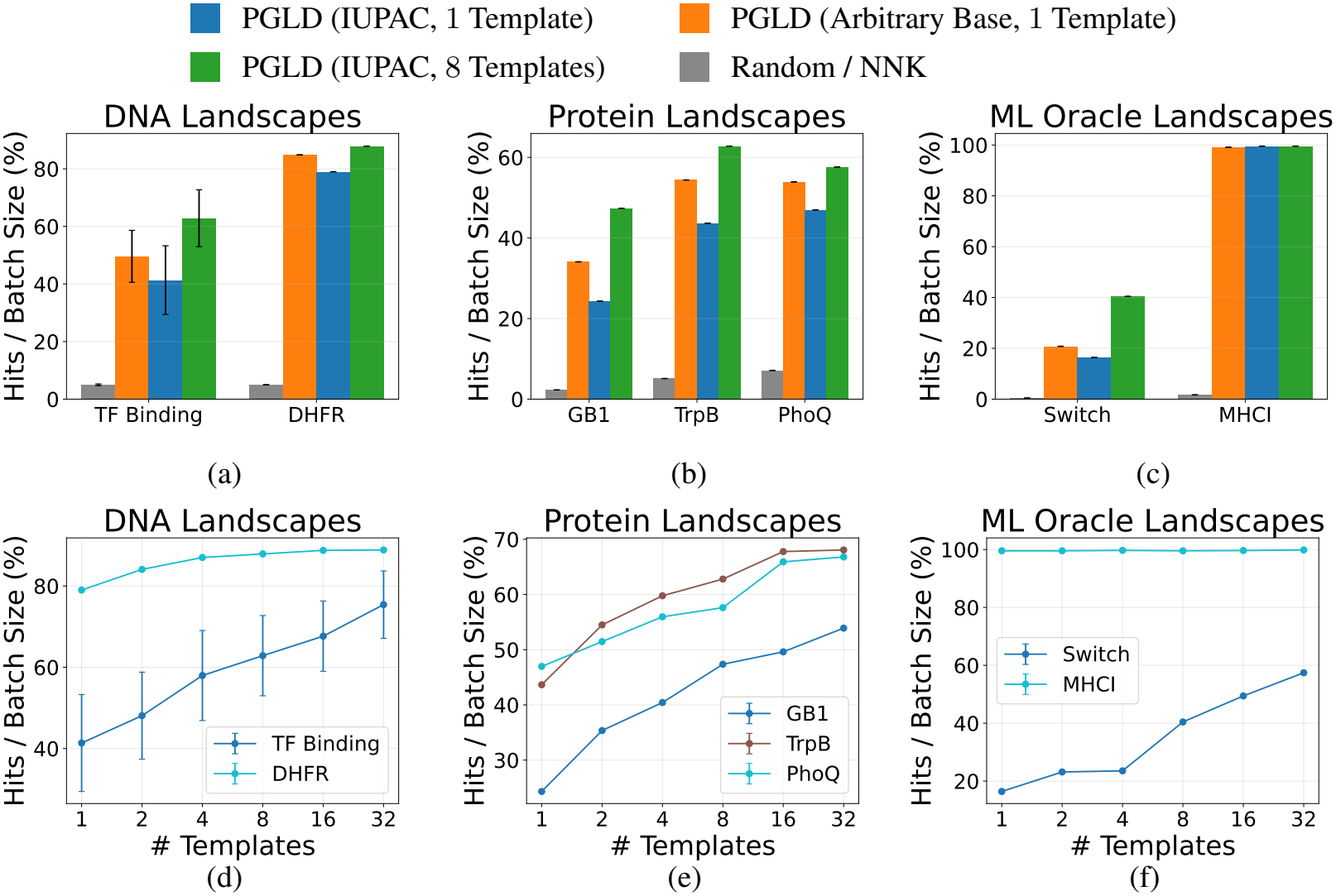
The PGLD optimizer, expected hits objective. **a**.-**c**. For 3 groups of landscapes, we evaluate PGLD on IUPAC and arbitrary base chemistries using the Expected Hits objective. We show the objective value achieved on each landscape after library optimization. The more flexible arbitrary base chemistry leads to higher objective values when comparing the performance of a single template, however increasing to just 8 IUPAC templates leads to the highest optima across all landscapes. **d**.-**f**. We repeat the optimization but sweep the number of templates available to the IUPAC optimizer. We observe considerable improvements as the number of templates is increased.

In all experiments in this section, 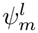 is always initialized N (0, 1) for all *m, l* when using PGLD with IUPAC bases. For arbitrary base chemistry, including the method of Zhu et al. [56], we have *L* × *M* mixed bases and we initialize their corresponding continuous parameters all in the same way (from N (0, 1)). In all experiments we use *N*_MC_ = 5 and a learning rate of 0.02.

In Table 5 we give the configurations used for each landscape. *ρ* is always set to either be the wild type fitness where one exists, or a sufficiently challenging threshold (e.g.for TF binding and MHCI). For Switch, we use the same threshold for a hit as was used in the original paper [55]. For the TF Binding landscape, we select 41 of these landscapes (all those indicated as a first round experiment), run PGLD and all baselines on all separately, and then report average results with standard deviation bars shown for variance in terms of the results across landscapes.

**Table 5:** Experimental settings for each landscape in Section 5.1.

| **Landscape** | **Batch Size** *B* | **Threshold** *ρ* | **Notes** |
|---|---|---|---|
| GB1 | 1,024 | 1.0 | – |
| PhoQ | 1,024 | 3.287 | – |
| TrpB | 1,024 | 0.092 | – |
| MHC | 2,048 | top 0.5% | Allele: HLA-A*02:01 |
| TF Binding | 256 | top 5% | Objective: average over 41 TFs |
| DHFR | 1,024 | 0.857 | – |
| Switch | 1,024 | 6.0 | Objective: ATP |

Full hyperparameters used to run PGLD on each landscape are given in Table 6. We find on some landscapes that for higher template counts more iterations were needed for the reward to converge, in which case we increase the number of iterations for larger template counts. The max queries per template per MC sample is the maximum number of times we query the landscape per MC sample for each template in the design. When an iteration contains fewer unique sequences per template than this number, we score all of them. For example, for landscapes with a batch size of 1024 and 256 max queries per template per MC sample, with 4 templates or more we query the landscape with all sequences, since the batch size is split among templates. Null means we always score the entire batch. We score protein sequences that have an early stop codon with a value lower than the fitness of any sequence on any landscape. For MHCI and TF Binding we use 0.0, for Switch −1.0 and for all others −20.0.

**Table 6:** Optimization parameters and settings for each landscape. The max queries per template per MC sample refers to the maximum number of unique sequences scored per template per MC sample, and when fewer unique sequences than this exist at a given iteration we score all of them.

| **Landscape** | **Batch Size** | **Base Iters** | **Iter Scaling** | **Max Queries/Template/MC sample** |
|---|---|---|---|---|
| GB1 | 1,024 | 10,000 | 2× at 16T, 4× at 32T | 256 |
| PhoQ | 1,024 | 10,000 | 2× at 16T, 4× at 32T | 256 |
| TrpB | 1,024 | 10,000 | 2× at 16T, 4× at 32T | 256 |
| MHCI | 2,048 | 5,000 | None | 256 |
| TF Binding | 256 | 25,000 | None | null |
| DHFR | 1,024 | 10,000 | 2× at 16T, 4× at 32T | 256 |
| Switch | 1,024 | 10,000 | 2× at 16T, 4× at 32T | 256 |

Theoretical diversity of a library is the number of unique sequences that can be possibly sampled by a given library design. For protein-scoring landscapes, this is the number of unique amino acid sequences. We do not include sequences containing a stop codon in the theoretical diversity count. For nucleotide-scoring landscapes this is the number of unique nucleotide sequences. Throughout the text, when computing theoretical diversity of libraries designed with arbitrary base synthesis, we mark amino acids or nucleotides at a given position as plausible to sample only if their probability of appearing at that position on that template exceeds 3%. This is approximately the probability of sampling the least common amino acid at a given position under an NNK design. In cases where we have a large number of templates, the Karp-Luby algorithm [21] is used to estimate the theoretical diversity of a library.

In Fig. 7a we show the fitness-diversity trade-off when varying *λ* with the Mean-Entropy objective. For a fixed *λ* and 32 templates, in Fig. 7c we show how the per-template allocation of credit leads to substantially steeper learning curves than a global credit assignment. We sweep across 5 learning rates for per-template and global, and plot the curve with the highest final reward value. The same sweep over learning rates was done for Fig. 2c. For the Mean-Entropy objective, all landscape and PGLD hyperparameters are the same as in Expected Hits experiments, besides the objective being swapped. Entropy is always estimated using the full batch size, while the mean fitness is estimated using only sequences that the landscape is queried with.

**Figure 7:**
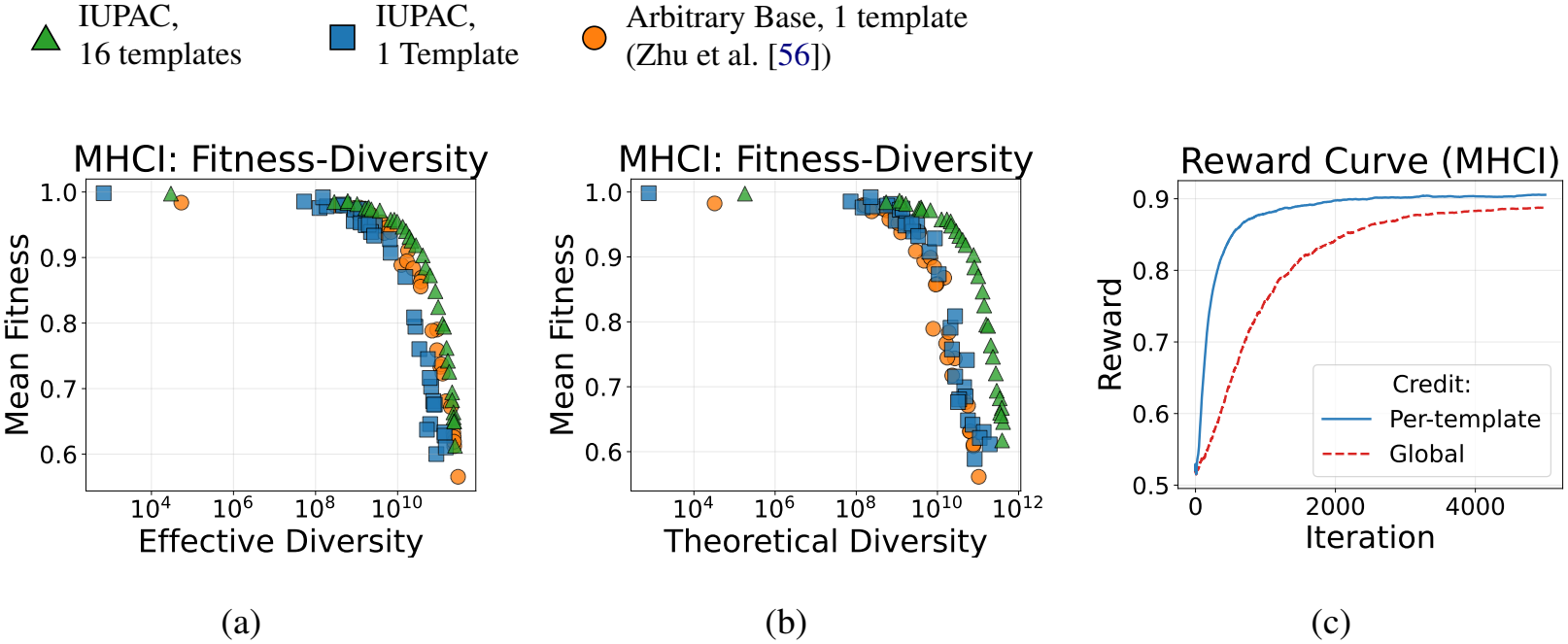
The PGLD optimizer, ablations. **a.** We repeat library optimization on the MHCI landscape with the mean-entropy objective, varying the value of *λ* to trade-off library diversity and fitness. Each point corresponds to a library trained with a different *λ* and different method. **b**. The same *λ* sweep, but on the x-axis we plot effective diversity (exponential of library entropy) instead of theoretical diversity. **c**. Reward curves for optimizing with PGLD, mean-entropy objective, on the MHCI landscape with 32 templates. Marginal gain credit assignment leads to sharper learning curves than global credit assignment.

## H Training with Limited Landscape Query Budget

When designing PGLD libraries, we seek to exhaust the throughput of our assay with the size of the final library *B*_final_. However, using a batch size that matches this throughput during optimization may not be feasible. For optimizing over fitness landscapes defined by an ML oracle as in Section 4.2 or Section 4.3, which are the method of choice for problems of practical relevance, querying the oracle is usually the rate-limiting step over sampling a batch of sufficient size. Therefore, we restrict ourselves to a limited budget of oracle queries *Q* per iteration of optimization. Here we investigate the number of hits in the final library with *B*_final_ = 10000 on the fully enumerated tetrapeptide landscapes (GB1, TrpB, PhoQ) with the IUPAC optimizer using 5 templates and *N*_MC_ = 5. To ensure similar difficulty of each task we choose the threshold for a hit as the 95th fitness percentile, yielding a total of 8000 possible hits per landscape. Additionally, we report results on the Influenza antibody landscape using *B*_final_ = 1000000 with 10 templates and *N*_MC_ = 10. The thresholds for hit sequences are set to the wildtype sequence fitness. We sample *Q* unique sequences and use the importance sampling estimator [23] to obtain an unbiased estimate of the batch reward. Experiments shown in this section use the Expected Hits objective introduced in Eq. (5), but examples of applying this idea to the Mean-Entropy objective can be found in Section 5.3 and Section 5.4. Fig. 8a demonstrates that overcoming computational constraints by reducing the batch size used in optimization *B*, and hence setting *B* = *Q*, generalizes poorly to the originally desired batch size. We observe that choosing *B* similar to the desired size of the final library and *Q << B* yields library performance comparable to *Q* = *B* at much reduced computation cost. The large desired library size *B*_final_ = 1000000 in the Influenza landscape does not allow for optimization with *Q* = *B*. Fig. 8b shows that we achieve a more than 100-fold increase in the number of predicted unique hits in the final library when scaling the batch size *B* = *B*_final_ but keeping the query budget *Q* constant.

**Figure 8:**
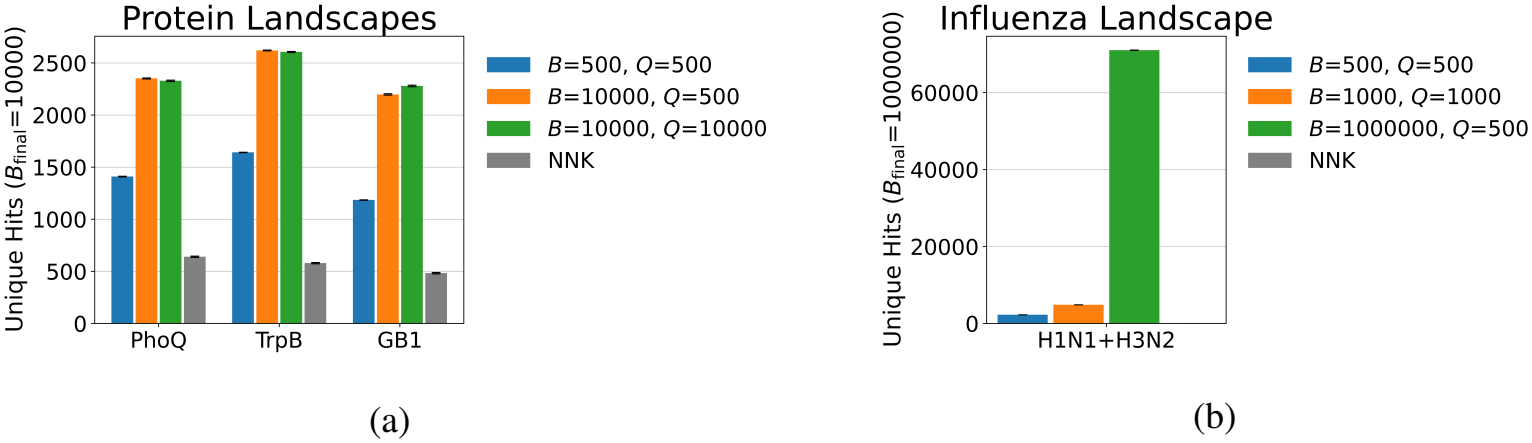
Generalizing to larger library sizes under a limited query budget. We plot the number of (predicted) unique hits in a large PGLD library of size *B*_final_ optimized with different batch size *B* and query budget *Q* on **a**. Enumerated protein landscapes and **b**. the C05 landscape based on DDMS data in Section 5.4. We report the average number of unique hits over 10 sampled final libraries.

## I Lab-in-the-loop

In this section, we provide implementation details for the experiments discussed in Section 5.2. We provide details on surrogate model training, baseline implementation, and parameters used.

In Fig. 9 we show the results from Fig. 3 of Section 5.2. We provide additional analysis on the distribution of sequences sampled by each algorithm, for each landscape, in Table 7. We demonstrate that even though PGLD was trained with a set hit threshold *ρ*, it is able to generally concentrate sequences at the top of the fitness distribution. We calculate the number of unique sequences above each of the following quantiles: 99.9, 99, 95, 90, and 85 sampled through all experiment iterations and show that PGLD yields at least a 40% increase in hits compared to other methods for 4 out of 5 small (*<* 10^6^ sequences) landscapes, no matter what threshold is used to define a hit. As an ablation study, we repeat the experiments of Section 5.2 but with varying number of templates for the PGLD library. The results can be found in Fig. 10 and demonstrate how PGLD performance improves with increasing template count.

**Table 7:** Unique sequence counts above quantile thresholds (averaged)

| Landscape | Method | ≥ 99.9% | ≥ 99% | ≥ 95% | ≥ 90% | ≥ 85% |
|---|---|---|---|---|---|---|
| DHFR | PGLD | **211.1** | **2146.9** | **5336.9** | 5773.7 | 6053.2 |
|  | Rational | 153.1 | 1627.0 | 5063.5 | **5910.4** | **6216.4** |
|  | Random | 31.1 | 282.8 | 1414.5 | 2835.3 | 4255.0 |
|  | % Incr | 37.9% | 32.0% | 5.4% | -2.3% | -2.6% |
| GB1 | PGLD | **140.9** | **1220.2** | **3870.9** | **5034.3** | **5615.2** |
|  | Rational | 76.1 | 661.3 | 2191.1 | 3200.0 | 3889.6 |
|  | Random | 26.9 | 275.4 | 1294.4 | 2378.4 | 3338.7 |
|  | % Incr | 85.2% | 84.5% | 76.7% | 57.3% | 44.4% |
| TF-binding | PGLD | **52.2** | **462.5** | **1688.0** | **2452.2** | **2801.6** |
|  | Rational | 21.0 | 173.2 | 775.9 | 1384.7 | 1976.6 |
|  | Random | 9.7 | 106.1 | 544.2 | 1094.8 | 1651.4 |
|  | % Incr | 148.6% | 167.0% | 117.6% | 77.1% | 41.7% |
| PhoQ | PGLD | **71.3** | **926.0** | **4088.4** | **5748.8** | **6821.3** |
|  | Rational | 22.7 | 482.3 | 2504.0 | 3801.1 | 4775.2 |
|  | Random | 15.2 | 285.1 | 1672.6 | 3167.9 | 4397.3 |
|  | % Incr | 214.1% | 92.0% | 63.3% | 51.2% | 42.8% |
| TrpB | PGLD | **153.7** | **1352.7** | **4686.6** | **5784.7** | **6271.7** |
|  | Rational | 87.6 | 634.9 | 1440.5 | 1712.5 | 1934.1 |
|  | Random | 44.1 | 389.6 | 1512.3 | 2431.1 | 3293.7 |
|  | % Incr | 75.5% | 113.1% | 225.3% | 237.8% | 224.3% |

**Figure 9:**
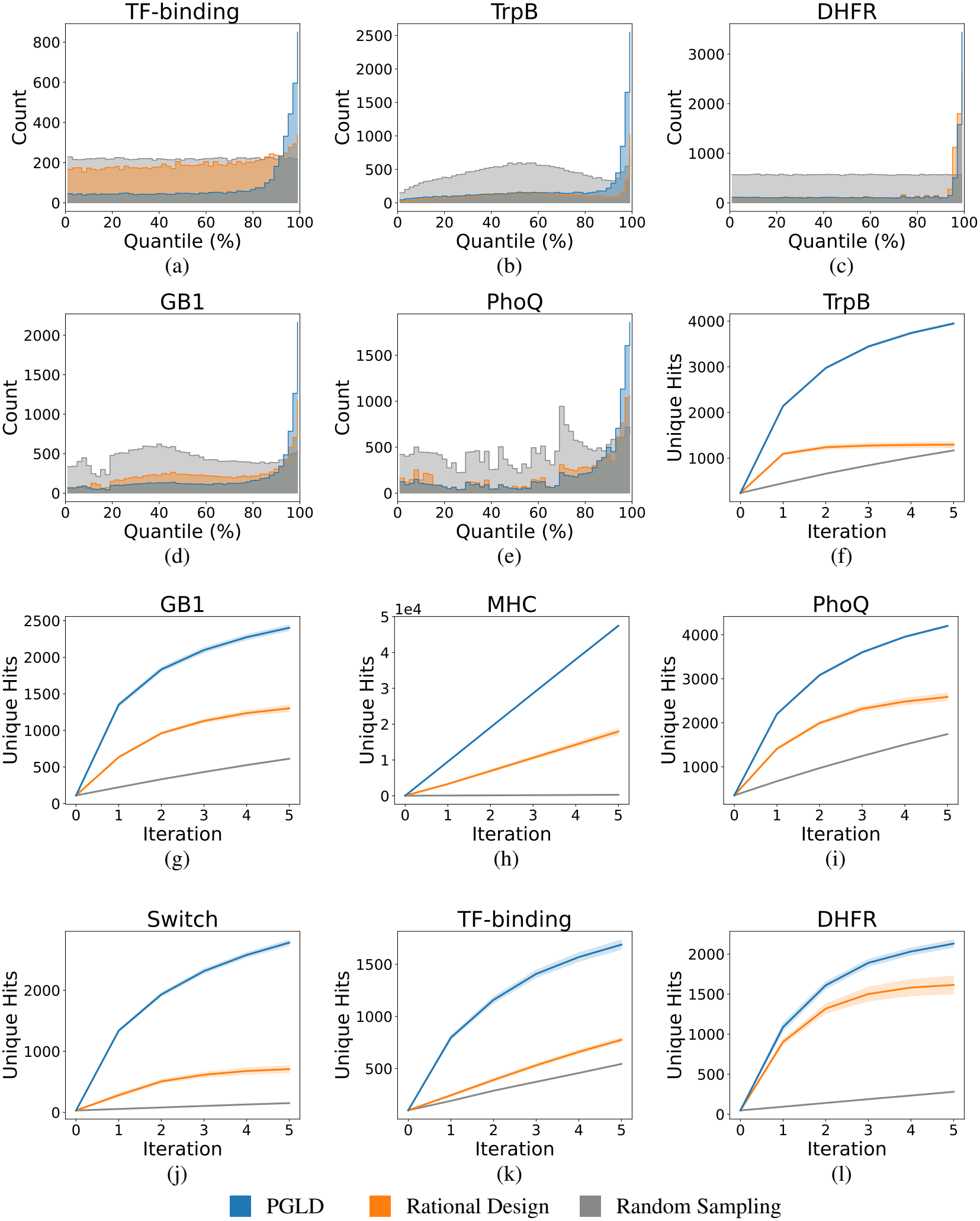
In-silico lab-in-the-loop experiments. For each of the seven landscapes described in Appendix F we provide comparisons of Algorithm 3, rational design and random sampling. All experiments consisted of 6 rounds. Round 0 is a random sampling round shared by all methods. We repeat the in-silico experiment ten times with different seeds. **a**.-**e.** For the five landscapes based on querying real data, we compare the quality of sequences sampled by each of the three methods. We show the quantile distribution of all unique, valid sequences sampled. All continuous measurements in the ground-truth datasets were mapped to quantiles for visualization. The results shown are averaged across ten random seeds. **f**.-**l**. For each landscape and method, we provide comparisons of the cumulative number of unique hits sampled at the end of each in-silico round of experimentation. We show the mean and the standard error across ten repeats with different seeds.

**Figure 10:**
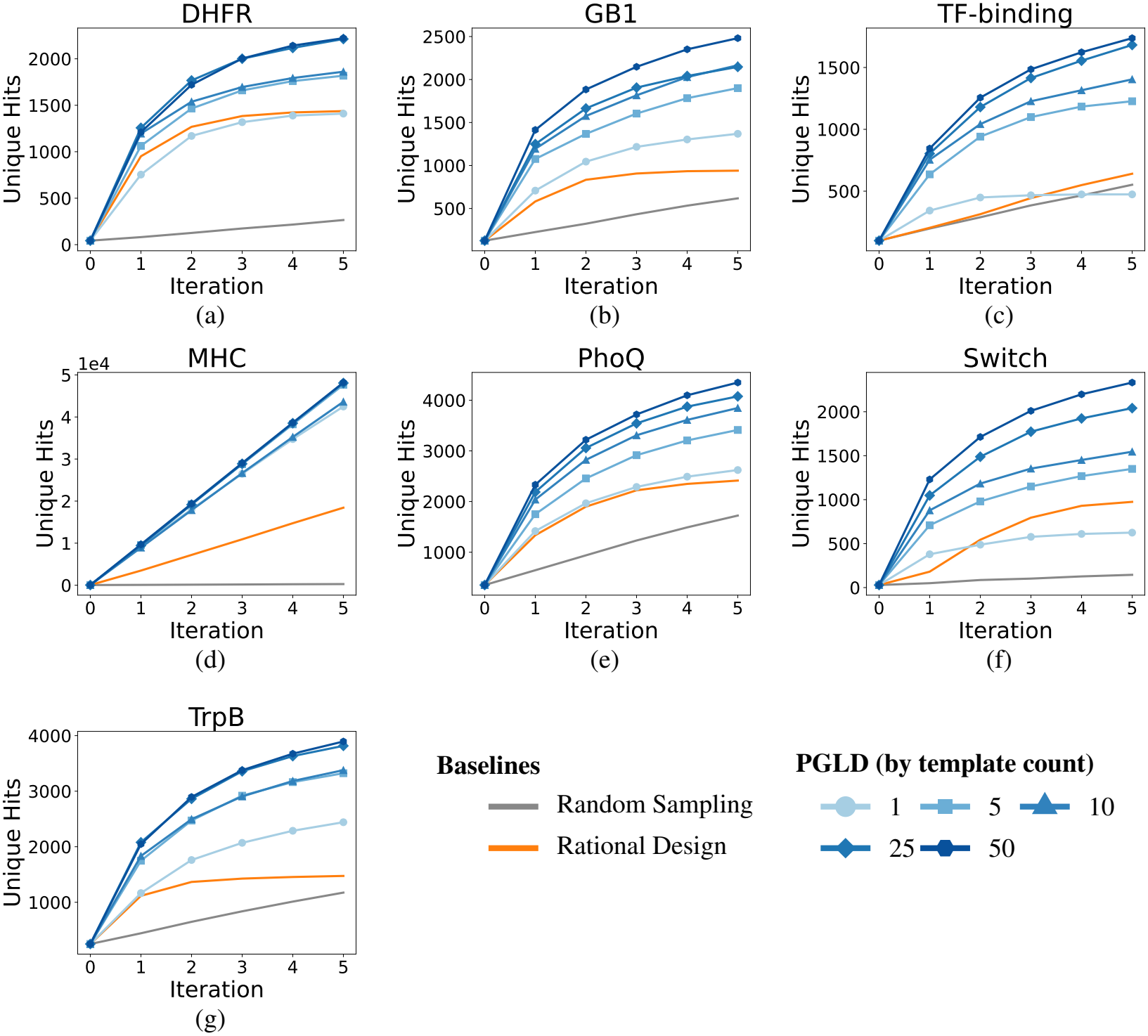
Template count ablation for in-silico lab-in-the-loop experiments. For each of the seven landscapes, we compare PGLD performance with varying numbers of templates (1, 5, 10, 25, 50) against rational design and random sampling baselines. All experiments consisted of 6 rounds, with round 0 being a random sampling round shared by all methods. **a**.–**g**. Cumulative number of unique hits sampled at the end of each in-silico round.

### I.1 Surrogate model training

The surrogate model used for all seven landscapes is a deep convolutional neural network. The input to the network is a one-hot encoded sequence where the encoding class is either one of the four nucleotides or one of the 20 amino acids. In the case of landscapes with DNA scoring the input has dimension *L* × 4 where *L* is the length of the sequence and in the case of landscapes with AA scoring the input has dimension *L* × 20. The model is trained with MSE loss on normalized targets to predict the target function for each landscape. Before training, the training dataset is deduplicated to include each unique sequence only once. For certain landscapes with skewed distributions, the target function is log-transformed before training. We provide the full architecture details in Table 8. The input to the model is shape *L* × *C* where *C* ∈ {20, 4}.

**Table 8:**
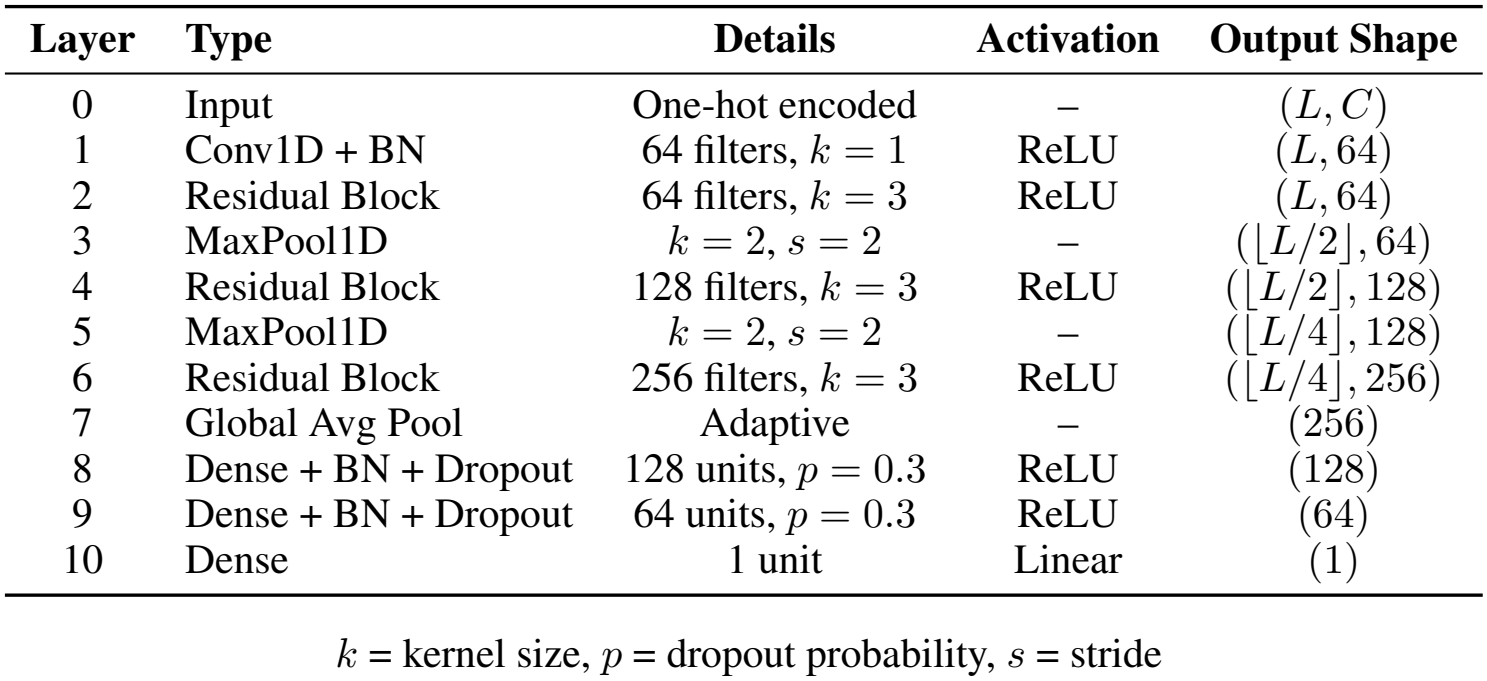
Surrogate model architecture.

| Layer | Type | Details | Activation | Output Shape |
|---|---|---|---|---|
| 0 | Input | One-hot encoded | – | (*L*, *C*) |
| 1 | Conv1D + BN | 64 filters, *k* = 1 | ReLU | (*L*, 64) |
| 2 | Residual Block | 64 filters, *k* = 3 | ReLU | (*L*, 64) |
| 3 | MaxPool1D | *k* = 2, *s* = 2 | – | (⌊*L*/2⌋, 64) |
| 4 | Residual Block | 128 filters, *k* = 3 | ReLU | (⌊*L*/2⌋, 128) |
| 5 | MaxPool1D | *k* = 2, *s* = 2 | – | (⌊*L*/4⌋, 128) |
| 6 | Residual Block | 256 filters, *k* = 3 | ReLU | (⌊*L*/4⌋, 256) |
| 7 | Global Avg Pool | Adaptive | – | (256) |
| 8 | Dense + BN + Dropout | 128 units, *p* = 0.3 | ReLU | (128) |
| 9 | Dense + BN + Dropout | 64 units, *p* = 0.3 | ReLU | (64) |
| 10 | Dense | 1 unit | Linear | (1) |

*k* = kernel size, *p* = dropout probability, *s* = stride

One residual block contains: Conv1D → BN → ReLU → Dropout(*p* = 0.15) → Conv1D → BN → Add(skip) → ReLU. Skip connections use 1 × 1 convolution when input/output channels differ.

Table 9 provides details on the hyperparameters used.

**Table 9:** Surrogate model training hyperparameters.

| **Hyperparameter** | **Value** |
|---|---|
| Optimizer | Adam |
| Learning rate | 10<sup>−3</sup> |
| Loss function | MSE |
| Batch size | 64 |
| Max epochs | 200 |
| Early stopping patience | 50 |
| Validation split | 20% |
| Target normalization | Z-score |
| Weight initialization | He normal |

### I.2 Rational Design Baseline

In this section we provide implementation details for the rational design baseline we use as comparison for PGLD. We will describe the algorithm in the case of a landscape with AA scoring first. At the start of round *i* we design an IUPAC template *T* from which a new batch of sequences *S*_*i*_ is sampled for measurement. Designing the template goes as follows: first we compute the target frequencies *ℱ* from the observed set *H*_*i*−1_. Target frequency *ℱ*_*p*_ tells us, for position *p*, what is the desired distribution over 20 possible amino acids. For each amino acid *a* and position *p* we set its target frequency to be the frequency of *a* at position *p* among all unique hit sequences in the history *H*_*i*−1_. Once we have defined *ℱ* we design an IUPAC template *T* so that for each degenerate codon *c*_*p*_ in *T* the MSE between the target frequency and the frequency of amino acids induced by *c*_*p*_ (denoted as **g**_*c*_) is minimized. The step of matching the target frequencies with **g**_*c*_ by minimizing the MSE is described in [28] and used in [42] for combinatorial library design. An additional stop codon penalty is added. In our experiments we set *λ* = 5. In Algorithm 4, the full algorithm across *T* rounds is described

#### Algorithm 4

Rational Design

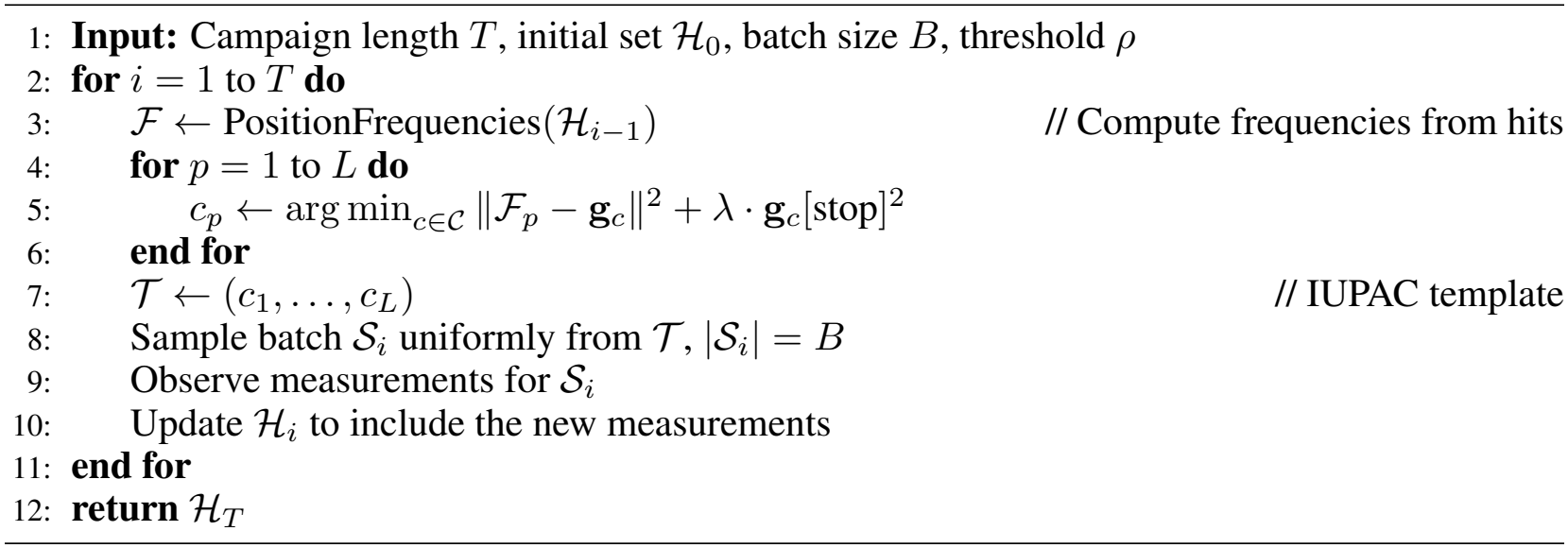

In the case of landscapes based on DNA scoring, the algorithm is simplified and instead of choosing codons which minimize the MSE with target amino acid frequencies, we can directly choose IUPAC bases which minimize the MSE between their induced nucleotide probabilities and the target nucleotide probabilities. The target nucleotide probabilities are set in the same way as the target amino acid probabilities - for each position *p* and nucleotide *n* we set its target frequency as the frequency of *n* at position *p* among all unique hit sequences.

### I.3 Landscape parameters

In Table 10, we provide details on relevant landscape related parameters. Batch sizes *B* were selected relative to the sizes of the landscapes. For some landscapes, their fitness value *f* was log-transformed before training the surrogate models. For MHCI, in this setting the fitness is − log_10_ of the MHCflurry predicted affinity in nM, rather than the affinity percentile used in Section 5.1.

**Table 10:** Experimental settings for each landscape in Section 5.2.

| Landscape | Batch Size *B* | Threshold ρ | Transformation | Notes |
|---|---|---|---|---|
| GB1 | 5,000 | 1.0 | log<sub>10</sub>(1 + *f*) | – |
| PhoQ | 5,000 | 3.287 | log<sub>10</sub>(1 + *f*) | – |
| TrpB | 5,000 | 0.092 | None | – |
| MHC | 10,000 | top 0.5% | −log<sub>10</sub>(*f*) | Allele: HLA-A*02:01 |
| TF Binding | 2,000 | top 5% | None | Objective: SIX6_REF_R1 |
| DHFR | 5,000 | 2.5 | log<sub>10</sub>(1 + *f*) | – |
| Switch | 5,000 | 6.0 | log<sub>10</sub>(1 + *f*) | Objective: ATP |

**Table 11:** Validation set evaluation metrics for the H1N1 and H3N2 surrogate models, stratified by the number of mutations from the wild-type sequence.

| **Strain** | **Metric** | **0–2 mutations** | **3 mutations** | **4–6 mutations** |
|---|---|---|---|---|
| H1N1 | Pearson's *r* | 0.815 | - | - |
|  | AUC-ROC | 0.943 | 0.807 | 0.668 |
| H3N2 | Pearson's *r* | 0.908 | - | - |
|  | AUC-ROC | 0.971 | 0.913 | 0.739 |

We used the same thresholds *ρ* as in Section 5.1. The only exception is the DHFR landscape for which we increased the threshold to make the task more difficult. Additionally, for the landscapes based on AA scoring, DNA sequences which produced stop codons were not considered hits.

### I.4 PGLD parameters

For running PGLD, we always used a learning rate of 0.02, 8000 iterations, 50 templates. During training on the trained CNN oracle, the maximum queries per batch per template is set to 25 for MHCI and for all other landscapes it is set to 100. During data collection and evaluation on the real landscape, all sequences in the batch are used to query the landscape.

## J Sampling from Generative Models In-vitro

We plot additional metrics from Section 5.3 in Fig. 11.

**Figure 11:**
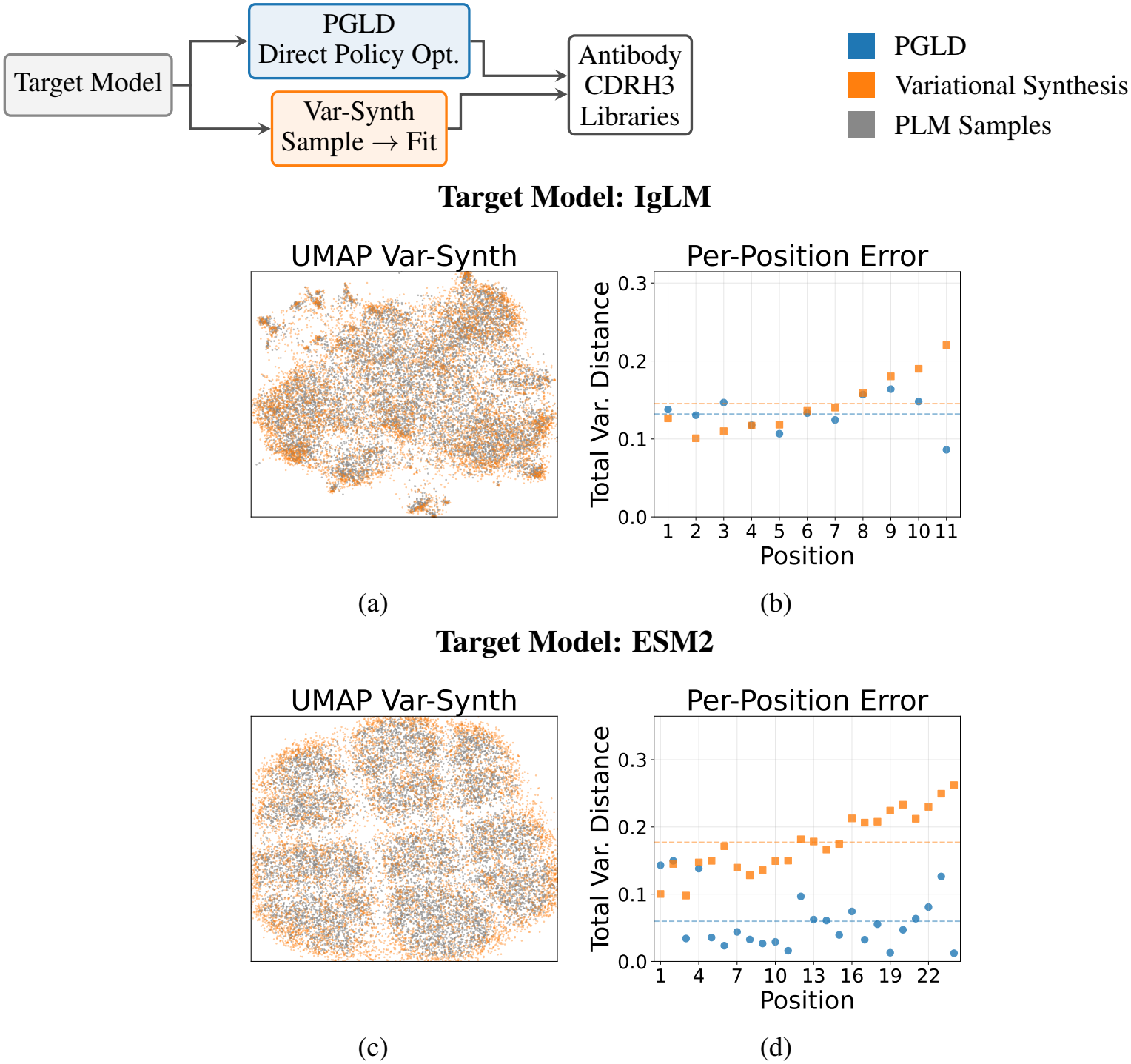
In-vitro sampling from generative models, additional visualizations. We compare sequences sampled from PGLD libraries and Variational Synthesis models, against the target PLMs. **(Top Row) Target Model: IgLM. a**. For Trastuzumab CDRH3s, Variational-Synthesis covers the target distribution in UMAP space. **b**. The total variational distance at each amino acid position between the PLM samples and each stochastic synthesis library, with the average as a dashed line. **(Bottom Row) Target Model: ESM2. c**. For C05 CDRH3s under a maximum edit distance of 6 from the wild type, Variational Synthesis covers the target distribution in UMAP space. **d**. The total variational distance at each amino acid position between the PLM samples and each stochastic synthesis library, with the average as a dashed line.

**Figure 12:**
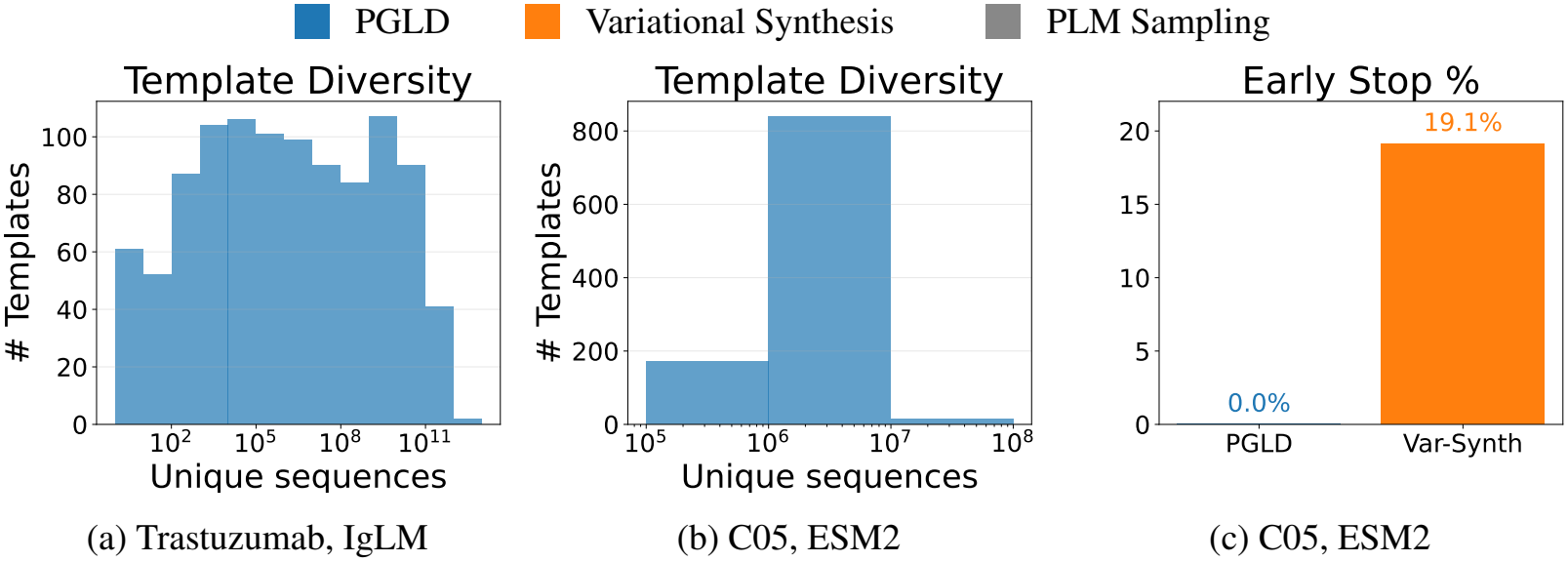
Sampling from a generative model supplemental figures. We plot per-template theoretical diversities for in-vitro generative model sampling experiments. For the 1024 templates in both of the PGLD-designed libraries in our two settings, we plot the theoretical diversity. For **a.** IgLM and **b**. ESM2. **c**. The proportion of sequences containing an early stop codon in the C05 libraries.

### J.1 Variational Synthesis Baseline

For running variational synthesis, we use the implementation released with Weinstein et al. [44] and always repeat with 3 different seeds, before selecting the model with lower perplexity for evaluating. For all hyperparameters, we use the defaults proposed in Weinstein et al. [44], so learning rate of 0.01, 5 Adam steps per EM update, 100 epochs (though convergence occurs significantly earlier), a minibatch size of 10000 and use Polyak-Ruppert averaging. We always train on 8 or 4 million sequences sampled from the PLM (described below) and use a separate 1 million sampled sequences in our evaluations. The 8 and 4 million sequences used for IgLM and ESM2 respectively is representative of the training data sizes used in Weinstein et al. [45] and we found that training on fewer (1 million) sequences for each task produced almost identical evaluation metrics. Since the designed sequences are short, we use a single pool (no assembly of shorter sequences into longer ones) as in Weinstein et al. [45].

### J.2 PLM Sampling and Scoring

IgLM is an autoregressive model trained specifically on antibody sequences [38], whereas ESM2 is a masked PLM trained on general protein sequences [25]. Because the 24-residue CDRH3 sequence of C05 is much longer than the average human CDRH3 at 11.6 (maximum length is 26) [50] and therefore maps to a low-density region of the IgLM distribution, we use ESM2 as the target PLM for C05 libraries.

For sampling sequences from IgLM, we condition on the [HUMAN] and [HEAVY] tokens, the left and right flanks of the full Trastuzumab heavy chain sequence, and use infill mode to sample CDRH3 sequences. We filter all sequences that don’t have a length of 11 (the length of the original CDRH3), and continue sampling until we reach 8.34 million samples. For scoring sequences with IgLM, both for creating Fig. 4a and for optimizing the PGLD library, we use the same conditioning and compute likelihoods of the specified CDRH3 sequence using infill mode.

For Gibbs sampling from ESM2, as a proposal distribution we sample a position *i* ∈ [*a, b*] uniformly, and then sample the amino acid at that position based on *p*(*s*_*i*_ | *s*_*/i*_). We then reject the proposal if it breaks the edit distance constraint, and otherwise we accept. We run 8192 parallel Gibbs sampling chains, each with a 1000 step burn-in, then saving sequences every 100 steps until we sample 4 million sequences. We tracked how many mutations each chain was accepting over 100 steps and found that this schedule produced sufficient mixing. As shown in Fig. 4f, the edit distance of most sampled sequences is 6. This is because there is no especially high likelihood assigned by ESM2 for being close to the C05 wild type, and there are significantly more sequences with edit distance 6 than e.g.edit distance 5. This Gibbs sampling is used to produce both the Variational Synthesis training data and the reference distributions used in evaluation.

Since exact likelihoods from a masked language model are not tractable to compute, we compute pseudolikelihoods for scoring sequences with ESM2. Given heavy chain sequence *s* with CDRH3 that starts at index *a* and ends at index *b*, we compute the pseudo log likelihood as

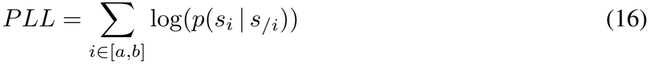

where *s*_*/i*_ corresponds to inputting the full sequence *s* with the *i*th position masked, and *p* is computed from the output logits of ESM2 at the *i*th position. For C05 with a 24 amino acid CDRH3, this corresponds to 24 forwards passes. We use this as the scoring mechanism both for PGLD library optimization and for scoring likelihoods in Fig. 4e. Note that the distribution induced by normalized pseudo-likelihoods will in general not correspond to the distribution of samples from the Gibbs sampler above. We alternate between them based on practicality (Gibbs for obtaining ESM2 samples, pseudo-likelihoods for scoring individual sequences). While PGLD is optimized against ESM2 pseudo-likelihood scores, the final library obtains reasonable correspondence to reference sequences sampled via Gibbs sampling.

### J.3 PGLD Hyperparameters and Initialization

To set *λ*, we run PGLD with smaller PLMs to define the target distribution (IgLM-S and ESM2-8M), and perform a small sweep over *λ* to find one that leads to a sufficiently large final theoretical diversity. We then run PGLD one time using the larger model and the chosen *λ*. The assumption made here is that smaller and larger versions of the same language model are sufficiently similar that the smaller model can be used for setting *λ*.

For final libraries, with IgLM we use *λ* = 1000 and for ESM2 we use *λ* = 3000. For both settings we use a PGLD learning rate of 0.02, sample a batch size of *B* = 2048 sequences per iteration (2 samples per template), MC samples *N*_mc_ = 1, and use marginal gain to assign credit to each template. For the likelihood assigned to constraint-breaking sequences, for IgLM we use *η* = −200, and for ESM2 we use *η* = −150. The values of *η* were set based on a small number of runs using smaller versions of each PLM. We score all 2048 samples each iteration for estimating the mean log likelihood, and all 2048 for obtaining a stochastic estimate of the entropy marginal gain per template.

In both settings, we initialize *ψ* by subsampling 1024 of the protein sequences used in the Variational Synthesis training data. We convert the amino acid sequence to a DNA sequence by inverting the codon table (if multiple codons produce the same amino acid, we select one randomly). Given 1024 DNA sequences, we initialize each template using each sequence. We set *ψ* such that for each position on each template, the probability of sampling the mixed base 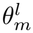 corresponding to that exact nucleotide is 0.995, and the remaining probability density is shared among the 14 other mixed bases.

### J.4 Evaluation Details

For evaluation, we sample 10^6^ samples from the PLM, 10^6^ from the PGLD library, and 10^6^ from the variational synthesis library. We subsample 10^4^ sequences from each of these for most plots: per position TVD, edit distances from wild type, early stopping rate, likelihood CDFs and UMAP visualizations. We subsample 10^5^ sequences from the PLM to train the projection for the UMAP plot. We subsample 10^5^ sequences from each library for the MMD [13] V-Stat computation, the BEAR score, and witness function score. Witness Score is the percentage of sampled sequences with score in the top 67% of MMD witness function scores among reference samples. BEAR score is the per-sequence Bayes Factor of the BEAR two-sample test. Effective diversity is the exponential of the entropy. For PGLD, we consider all sequences and for variational synthesis we consider both all sequences and the subset of valid-only sequences as variational synthesis has a non-negligible number of sequences that violate the edit distance and no-early-stop constraints. Effective diversity only considers no-early-stop sequences. For the per position TVD and likelihood CDF we include sequences that stopped early or had a constraint-breaking edit-distance (just scored as having 0 likelihood for the CDF plot). For the edit distance histogram and UMAP we don’t include early stopping sequences.

For the methodology of producing the TVD per-position plot, the UMAP visualizations, MMD test, witness scores, and BEAR scores, we follow the implementation described in Weinstein et al. [45]. For the UMAP visualizations, we train the projection on only reference sequences, while Weinstein et al. [45] used samples also from the Variational Synthesis library for training the projection.

Experiments to optimize a library with PGLD using IgLM were run on 1 Nvidia RTX 4090 for under 48 hours. Experiments to optimize a library with PGLD using ESM-150M were run on 8 Nvidia RTX 4090s for under 48 hours. For drawing training and testing samples from both language models, up to 8 Nvidia RTX 3090s were used for under 24 hours.

## K Influenza Antibody Model Fitting

### K.1 DDMS Data

To generate the DDMS dataset, we displayed the C05 antibody Fab-region variants on the surface of yeast cells and measured binding to the HA1 protein of the target strain using fluorescence activated cell sorting (FACS).

The positive and negative sorting gates were defined using the wild-type C05 Fab as positive control and the unstained library as negative control. Following a single round of enrichment (one round of cell sorting), both populations were analyzed via deep sequencing to obtain raw read counts.

The double deep mutational scan produces a separate dataset D for each target consisting of a list of tuples {*s, n*_*pos*_(*s*), *n*_*neg*_(*s*)}. For a sequence *s, n*_*pos*_(*s*) is the number of reads matching that sequence in the positive pool after screening with yeast display against the target, and *n*_*neg*_(*s*) is the number of reads in the negative pool. One can then compute the log enrichment of each sequence, a proxy for binding:

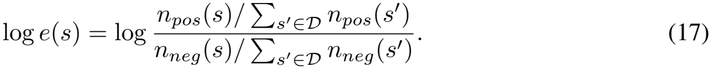

A small pseudocount is applied to positive and negative read counts to make them strictly positive.

### K.2 Objective Function

We train separate models for each target, but here for brevity we drop from the notation an explicit dependence on the target. Given a target and dataset D, we aim to train a model 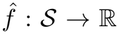 that predicts log enrichment from sequence. The DDMS dataset is only designed to contain mutations within 2 edits of the wild type C05 CDRH3, so by training a model 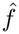, we can use it as the objective within PGLD to design libraries that produce sequences with higher edit distances predicted to bind the target.

One possibility is to train 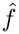 to directly predict log *e*(*s*) using, e.g., a mean-squared error loss. We instead framed the problem as a logistic regression task. For any given sequenced read, we model

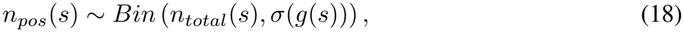

where *n*_*total*_ = *n*_*pos*_ + *n*_*neg*_ and *g* : S → ℝ is a function we learn, and *σ* is the sigmoid function.

We can then learn a neural network estimate of *g* by minimizing the Binary Cross-Entropy (BCE) loss, weighted by total observation count.

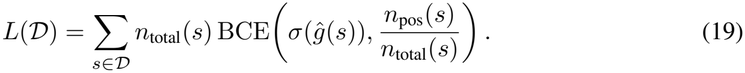

To prevent too much weight being placed on a single sequence, in practice we clip *n*_*total*_(*s*) in the equation above at 1000.

One benefit of this classification approach over regressing on log enrichment is that we can learn from *s* ∈ *D* where *n*_*total*_(*s*) is very small, making a log enrichment calculation for that sequence very noisy. In fact, we found that due to small synthesis or PCR errors in the construction of the DDMS library, the dataset also contained small numbers of reads for many triple and quadruple mutations. These unexpected sequences were unlikely to be from sequencing errors, since sequences with very low total counts were filtered out. The classification objective allowed us to also learn from these reads. Since *ĝ* estimates the log-odds of a read being positive, it is equivalent to a log enrichment estimate up to an additive constant. We therefore train *ĝ*_*H*1*N*1_ and *ĝ*_*H*3*N*2_, then use them as the surrogate fitness for each target.

### K.3 Model Architecture

Models were trained on a random train-val split of the DDMS data. The primary metrics for determining architecture choices were Pearson’s correlation coefficient between predictions from *ĝ* and log enrichments of validation set sequences, and AUC-ROC curves for classifying individual reads on the validation set.

We finetune an ESM2 model with an additional transformer head on top of the last layer. The full heavy chain of C05 is used for conditioning, but the added transformer head only conditions on embeddings at positions belonging to the 24 amino acid CDRH3.

We used the ESM2-8M model as the base for our final models. Scaling to larger model sizes did not show an improvement in terms of the two evaluation metrics we tracked. This may be because the DDMS dataset has a large number of unique sequences (over 10^5^), meaning that a strong base model is not necessary to achieve strong performance on the evaluations we tracked. It has been observed before that scaling base PLM model size does not correspond to improvements on downstream tasks [6, 18].

Both models utilized identical hyperparameters: the base PLM frozen, a batch size of 4096, a learning rate of 3 × 10^−4^, and a dropout rate of 0.1. The transformer head had depth 3, 8 attention heads and width 128. Training continued for a maximum of 300 epochs with an early stopping patience of 20 epochs. We experimented with varying the learning rate, unfreezing the base PLM weights, and varying the complexity of the added head, but these hyperparameters represent those used for training the final models. Models were trained on a single Nvidia RTX 3090 GPU for up to 90 minutes.

### K.4 Model Performance

We stratified our validation set by mutational distance from the wild type, and show our two primary metrics as measured on each group. These were Pearson’s correlation between predictions *ĝ* and log enrichments on the validation set, and AUC-ROC curves for classifying individual reads on the validation set.

Note that since all mutational distances higher than 2 were from synthesis errors or PCR errors, they represented relatively low read counts. Since for 3+ mutations almost all sequences had all positive or all negative reads, the computed log enrichment values were unreliable and hence we only show classification metrics. The AUC-ROC values on group 4 − 6 suggests that the trained models still carried some signal on mutational distances of up to 6 mutations. In some experimental runs (final models not used downstream), we trained similar models only on 1-mutation data from the DDMS and predicted 2-mutation log enrichments. We found Pearson’s r of 0.782 for H3N2 and 0.684 for H1N1.

## L Influenza Antibody Library Optimization

In the final PGLD library, we used *λ* = 1.25 and obtained a theoretical diversity of 9.78 × 10^5^. The surrogate score we use for early-stopping sequences is −20, consistent with our experiments in Section 5.1 and Section 5.2.

To initialize each template, we used a similar strategy to that in Appendix J to set each template’s *ψ*_*m*_ to have a 95% chance of sampling the wild-type nucleotide at each position, and a 5% chance of uniformly sampling any nucleotide at that position. We give the hyperparameters for the trained PGLD library in Table 12.

**Table 12:** PGLD C05 library hyperparameters.

| **Hyperparameter** | **Value** |
|---|---|
| Number of templates | 64 |
| Learning rate | 0.02 |
| Iterations | 12000 |
| Batch size | 10000 |
| MC samples | 10 |
| Landscape queries/template/MC sample | 2 |

So that it could be possible to later sequence and compute log enrichment values for each sequence from the PGLD library, we duplicated some templates to achieve a more uniform distribution over sequences. We assumed that the assay throughput was roughly 10^8^ sequences. We duplicated templates a number of times proportional to their theoretical diversity. Let the theoretical diversity of the *m*-th template be *d*_*m*_. We included a total of ⌈*d*_*m*_*/*21000⌉ copies of that template, so for any given original template, the average number of reads a sequence gets was at least ∼ 10^8^*/*(101 × 21000) ≈ 47. The exact value of 21000 was chosen as a trade-off between the amount of template duplication and the library uniformity. After duplicating several templates, the final library had 101 templates. All in-silico evaluation of the library was performed with the original 64 template design, and the experimental validation was performed with the 101 template design containing duplicates.

The Rational Design library was designed according to the general algorithm in Algorithm 4 but with several additional heuristics applied for real-world use. First, since yeast display of the DDMS library detects positives and negatives per-read, not per-sequence like our experiments in Section 4.2, the target distribution for each amino acid is set to be the distribution of reads at that position in the H1N1 and H3N2 DDMS positively binding libraries pooled together. A threshold is used to set the target probability of very low frequency amino acids to 0. The 5 positions with the highest mutation frequency in the positive pool were sub-selected to be varied to match the target distribution, and all other positions were kept fixed to the wild type amino acid. Focusing on a most permissive subset of amino acids to mutate allows for controlling the library theoretical diversity.

When evaluating libraries in-silico for expected unique hits (Fig. 5), we define a hit as a sequence whose predicted log enrichment under both *f*_*H*1*N*1_ and *f*_*H*3*N*2_ exceeds the ground-truth log enrichment of the wild-type C05 sequence measured in the DDMS dataset. We explicitly use the empirical DDMS values as the threshold rather than the surrogate models’ predictions for the wild-type sequence. This is because the surrogate models tend to overestimate the wild-type performance, both in terms of raw score and in terms of what proportion of double mutations in the DDMS data had higher log enrichment than the wild type.

We scored the final designed PGLD, Rational Design and DDMS libraries using the *f*_*H*1*N*1_ and *f*_*H*3*N*2_ models on a sample of 10000 sequences from each library. Figure 13a and Fig. 13b show the predicted fitness scores for each target, and their sum, which is used as reward during optimization, is depicted in Fig. 13c. We found that the PGLD library tended towards very high-scoring sequences, as expected since it was directly optimized on these objectives, while the other two libraries were not. Notably, Fig. 13d demonstrates that PGLD library sequences achieve high predicted fitness scores under both models simultaneously. We did not impose a particular edit distance constraint on the PGLD library as we did in Section 5.3.

**Figure 13:**
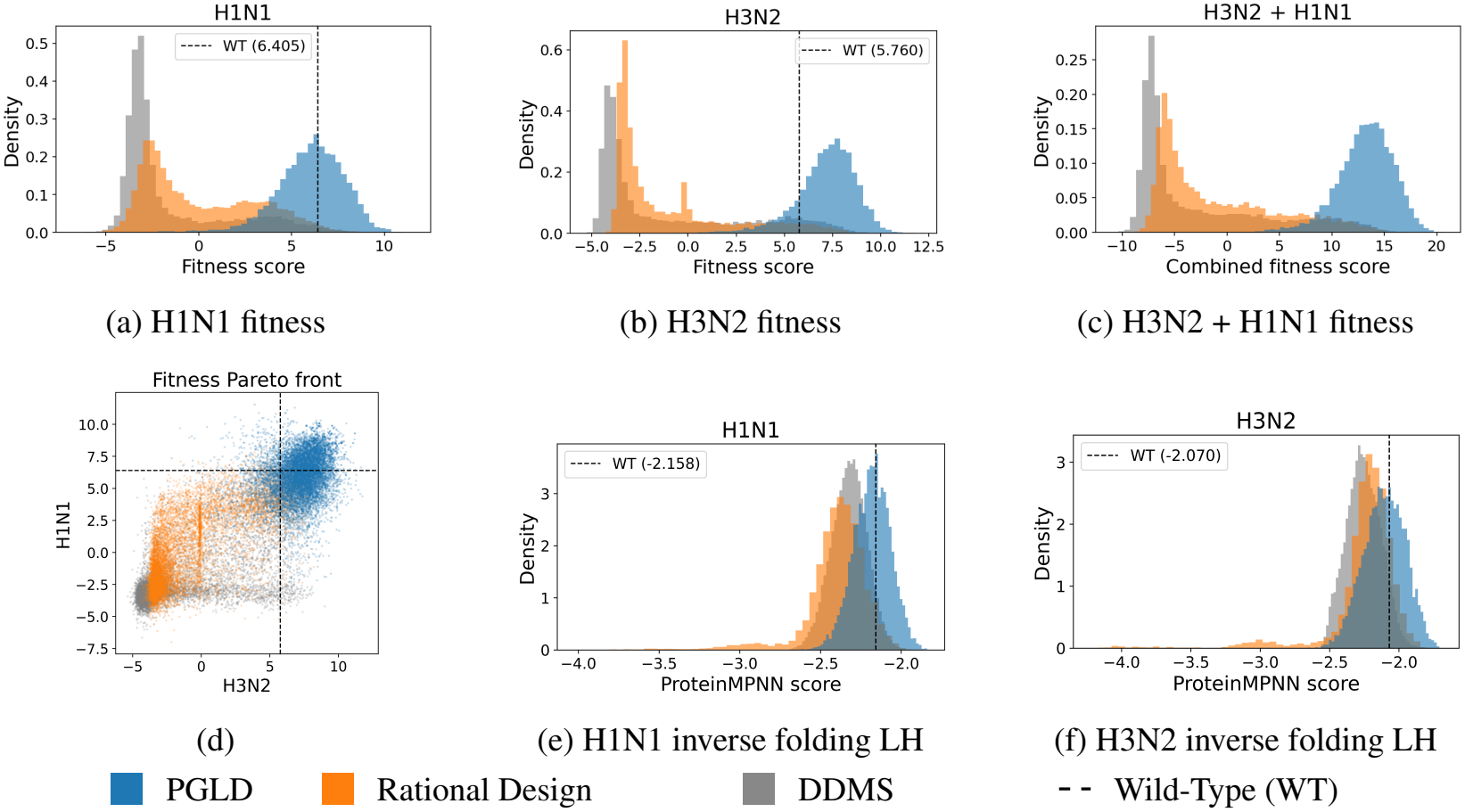
C05 antibody engineering, additional library visualizations. We plot the empirical distribution of the predicted fitness evaluated on 10000 sequences sampled from each library with models **a**. *f*_H1N1_, **b**. *f*_H3N2_, and **c**. their sum *f*_H1N1_ + *f*_H3N2_. **d**. visualizes the fitness values from a and b in a 2D-scatterplot. We also show the empirical distribution of ProteinMPNN inverse folding likelihoods when conditioning on Boltz2-generated structures of the C05 wild-type antibody in complex with the **e**. H1N1 and **f**. H3N2 antigen.

We scored the designed library in-silico based on an inverse folding model (ProteinMPNN) [8]. For both the H1N1 and H3N2 strains, we co-folded the sequence with the C05 wild type using Boltz2 (neither of these complexes can be found in the PDB) [33]. We used the solved C05 wild-type structure from entry 4FP8 as the template, and the solved antigen structure (different strain) from the 4FP8 complex for the antigen template. For each strain, we chose a final complex structure by sampling 50 candidate structures from Boltz2, then selecting the one with highest iPTM score (for H3N2: 0.77 iPTM) and verifying that the structure looked plausible. We conditioned on the chosen structure and the entire C05 sequence except the CDRH3 in MPNN, and used it to compute sequence likelihoods.

To validate MPNN likelihoods as a proxy for sequence fitness, we computed the MPNN likelihood of all double mutations in the H3N2 DDMS dataset and found that the MPNN likelihoods had a Pearson’s Correlation Coefficient of 0.30 (0.28 for H1N1). This suggests that the MPNN likelihoods conditioned on structure hold some information about what sequences are likely to have a higher log enrichment. However, we found that MPNN likelihoods had no selectivity based on the target: when swapping the target structures for the H3N2 and H1N1 strain we noticed no degradation in correlation with the DDMS log enrichment values.

We sampled 10000 sequences from the DDMS, Rational Design, and PGLD libraries and show their MPNN likelihood distribution for each target in Fig. 13e and Fig. 13f. We found that in the PGLD library sequences generally scored higher likelihoods under MPNN, suggesting they are more likely given the target complex structure.

Experiments to optimize a C05 library with PGLD were run on 1 Nvidia RTX 3090 and took approximately 13 hours.

The PGLD library was ordered via the IDT OligoPools product. The Rational Design library was ordered as an IDT Ultramer. The original DDMS library was ordered with IDT OligoPools. For validating that libraries corresponded to the ordered designs, we subsampled from the library after amplification in E.coli and performed Sanger sequencing on picked colonies. We could then verify that all sequences picked were possible under the ordered library design.

After performing fluorescence activated cell-sorting (FACS), we sequence the pre-sort population, post-sort positives and post-sort negatives for all library-antigen combinations. The PGLD sort and subsequent sequencing is performed twice. A visualization of the experimental workflow in yeast display is shown in Fig. 14. A binder is, for the purpose of this study, defined as any amino acid sequence that is feasible under the original library design, with ≥ 40 pre-sort reads, and a positive vs negative log ratio greater than the wild type’s. Raw binder counts in the sequencing data are not meaningful to compare across libraries due to differing read-depths, batch effects and within-library binding competition. Therefore our focus with the sequencing data is studying the diversity and mutational distances of classified binders for each library.

**Figure 14:**
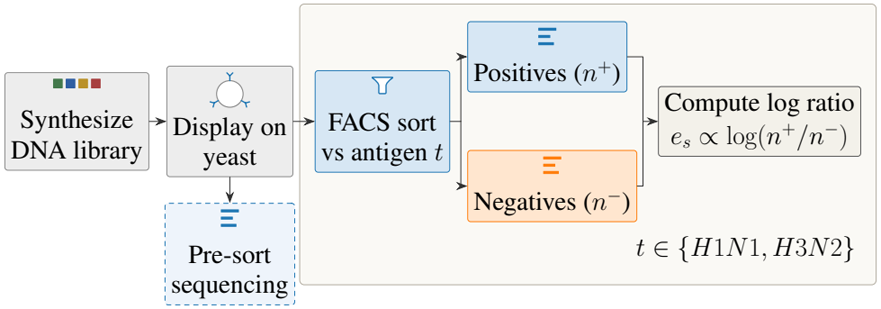
The experimental workflow in yeast display: after displaying on yeast, sort for binding against each antigen with fluorescence-activated cell sorting (FACS) and use sequencing of the positive and negative pool to compute a log-ratio score for each sequence.

For analysing sequence-level data from sorted positive and negative populations, as in Fig. 5d and Fig. 5e, we restrict the analysis to sequences intended under the original library design. This is to eliminate sequences that may have occurred due to synthesis or sequencing errors. Log ratio scores are computed for every sequence with a minimum pre-sort read depth (≥ 40 unless otherwise stated) against each antigen, as in Eq. (17). All sequences with log ratio score greater than the wild type’s score on the same screen are classified as a binder. For cases where we sorted twice (PGLD) we merge together sequencing data from both sorts.

On a sequence-by-sequence level, binders are not directly comparable across screens. This may be due to batch level effects but also competition effects within libraries. For example, when taking sequences appearing in both the Rational Design and PGLD libraries, we found that almost 100% of these sequences were defined as binders under the Rational Design sequencing data, while the PGLD data showed a mix of binders and non-binders (Fig. 15a).

**Figure 15:**
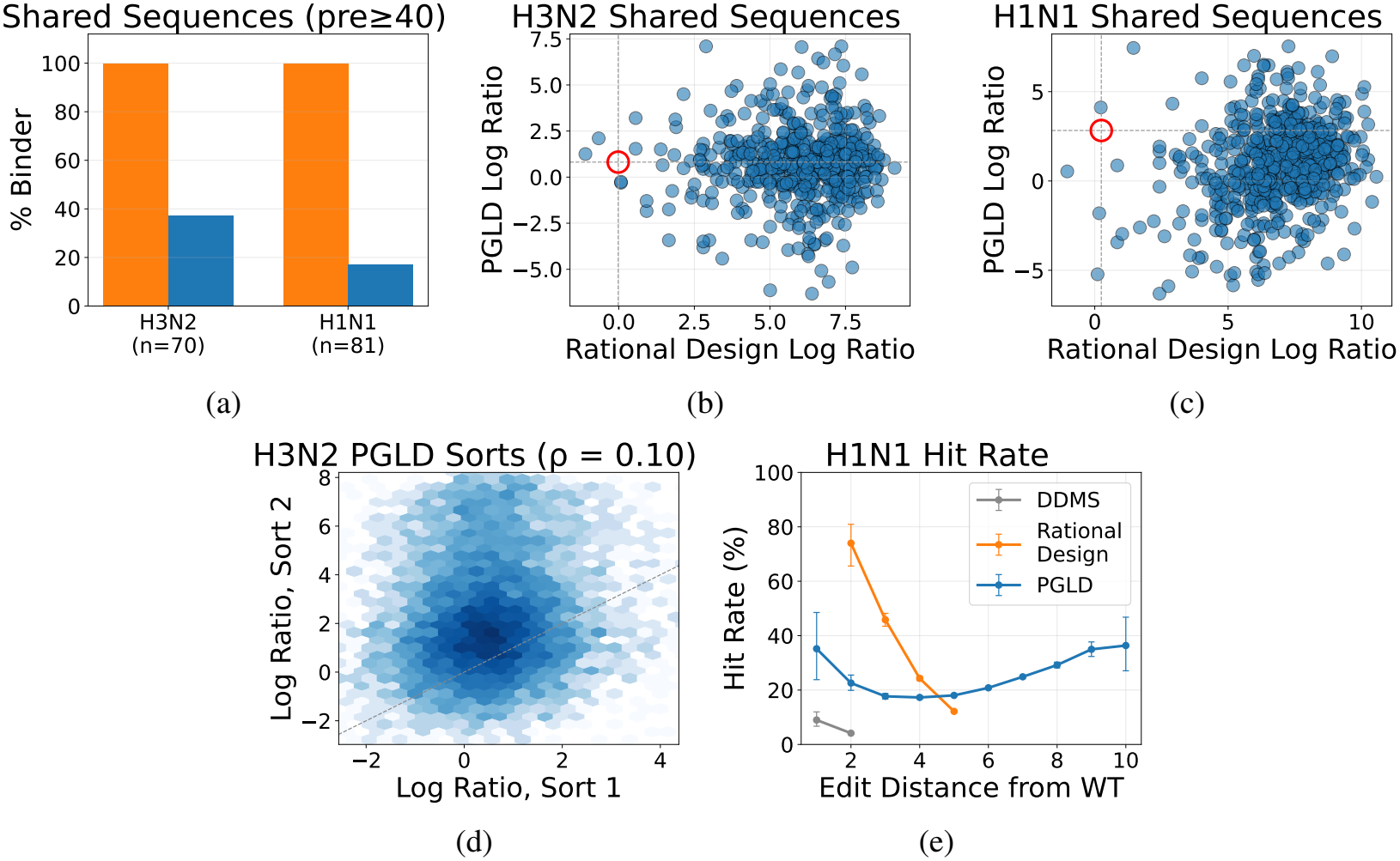
C05 antibody engineering, sequence-level supplementary results. **a.** For sequences with ≥ 40 pre-sort reads in both the Rational Design and PGLD sequencing data, all are classified as binders under the Rational Design data, but less than half are in the PGLD data. **b, c**. Log ratio scores for sequences (pre-sort reads ≥ 20) in PGLD and Rational Design sequencing data. The red circle marks the wild type. **d**. There is low Spearman’s correlation in log ratio scores for sequences with ≥ 40 pre-sort reads on both H3N2 PGLD library sorts. **e**. The proportion of sequences with ≥ 40 pre-sort reads classified as binders at each edit distance: PGLD shows stable hit rates with increasing edit distance. Error bars show 95% Wilson score confidence intervals.

We find that the very high proportion of binders for H3N2 in the PGLD library (Fig. 5c) leads to weak selection during a single round of yeast display. 100% of sequences appearing in both the H3N2 PGLD and Rational Design data are classified as binders in the Rational Design data (Fig. 15a). We find low reproducibility between the repeat sorts for the PGLD library on H3N2 (Fig. 15d). We hypothesize that this is due to the very low truly negative binding population resulting in yeast display imposing very weak selection. For H1N1 we find strong consistency across the repeat sorts for the PGLD library (Fig. 5d), and we find that this consistency increases at higher pre-sort read depths. We find a Spearman’s correlation of 0.88, at pre-sort minimum read depth of 100, compared to 0.80 at read depth ≥ 40. Therefore, we focus analysis on sequences defined as binders for the H1N1 target in each library. We find that the hit rate at each edit distance decreases with increasing edit distance for the Rational Design library while staying stable for the PGLD library (Fig. 15e).

## References

[1] Agarwal, A. A., Harrang, J., Noble, D., McGowan, K. L., Lange, A. W., Engelhart, E., Lahman, M. C., Adamo, J., Yu, X., Serang, O., et al. (2025). AlphaBind, a domain-specific model to predict and optimize antibody–antigen binding affinity. In MAbs, volume 17. Taylor & Francis.

[2] Barrera, L. A., Vedenko, A., Kurland, J. V., Rogers, J. M., Gisselbrecht, S. S., Rossin, E. J., Woodard, J., Mariani, L., Kock, K. H., Inukai, S., et al. (2016). Survey of variation in human transcription factors reveals prevalent DNA binding changes. Science, 351(6280):1450–1454.

[3] Bishop, C. M. and Nasrabadi, N. M. (2006). Pattern recognition and machine learning, volume 4. Springer.

[4] Boder, E. T. and Wittrup, K. D. (1997). Yeast surface display for screening combinatorial polypeptide libraries. Nature biotechnology, 15(6):553–557.

[5] Brixi, G., Durrant, M. G., Ku, J., Naghipourfar, M., Poli, M., Sun, G., Brockman, G., Chang, D., Fanton, A., Gonzalez, G. A., et al. (2026). Genome modelling and design across all domains of life with Evo 2. Nature, pages 1–13.

[6] Catrina, D., Bepler, C., Sledzieski, S., and Singh, R. (2026). Reverse distillation: Consistently scaling protein language model representations. arXiv preprint arXiv:2603.07710.

[7] Cornish-Bowden, A. (1985). Nomenclature for incompletely specified bases in nucleic acid sequences: recommendations 1984. Nucleic acids research, 13(9):3021.

[8] Dauparas, J., Anishchenko, I., Bennett, N., Bai, H., Ragotte, R. J., Milles, L. F., Wicky, B. I., Courbet, A., de Haas, R. J., Bethel, N., et al. (2022). Robust deep learning–based protein sequence design using ProteinMPNN. Science, 378(6615):49–56.

[9] Desautels, T., Krause, A., and Burdick, J. W. (2014). Parallelizing exploration-exploitation tradeoffs in Gaussian process bandit optimization. The Journal of Machine Learning Research, 15(1):3873–3923.

[10] Didi, K., Reidenbach, D., Penner, M., Ravi, S., Case, M., Nichols, M., Swanson, E., Reis, A., Prescott, M., Qian, Y., Qian, D., Yang, J., Li, W., Li, L., Shonai, D., Gay, S., Mallik, B. B., Chim, H. Y., Chen, L., Juantay, M. A., Klein, H., Macintyre, A., Secor, M., Granata, D., Cao, Z., Zhou, G., Geffner, T., Chen, X., Livne, M., Zhang, Z., Zhang, T., Bronstein, M. M., Steinegger, M., Deibler, K., Soderling, S., Khmelinskaia, A., Hollfelder, F., Dallago, C., Kucukbenli, E., Vahdat, A., Ogden, P., and Kreis, K. (2026). Latent generative search unlocks de novo design of untapped biomolecular interactions at scale. https://research.nvidia.com/labs/genair/proteina-complexa/assets/proteina_complexa_validation.pdf.

[11] Ekiert, D. C., Kashyap, A. K., Steel, J., Rubrum, A., Bhabha, G., Khayat, R., Lee, J. H., Dillon, M. A., O’Neil, R. E., Faynboym, A. M., et al. (2012). Cross-neutralization of influenza A viruses mediated by a single antibody loop. Nature, 489(7417):526–532.

[12] Fowler, D. M. and Fields, S. (2014). Deep mutational scanning: a new style of protein science. Nature methods, 11(8):801–807.

[13] Gretton, A., Borgwardt, K. M., Rasch, M. J., Schölkopf, B., and Smola, A. (2012). A kernel two-sample test. The journal of machine learning research, 13(1):723–773.

[14] Harris, D. T., Wang, N., Riley, T. P., Anderson, S. D., Singh, N. K., Procko, E., Baker, B. M., and Kranz, D. M. (2016). Deep mutational scans as a guide to engineering high affinity T cell receptor interactions with peptide-bound major histocompatibility complex. Journal of Biological Chemistry, 291(47):24566–24578.

[15] Hayes, T., Rao, R., Akin, H., Sofroniew, N. J., Oktay, D., Lin, Z., Verkuil, R., Tran, V. Q., Deaton, J., Wiggert, M., et al. (2025). Simulating 500 million years of evolution with a language model. Science, 387(6736):850–858.

[16] Hie, B. L., Shanker, V. R., Xu, D., Bruun, T. U., Weidenbacher, P. A., Tang, S., Wu, W., Pak, J. E., and Kim, P. S. (2024). Efficient evolution of human antibodies from general protein language models. Nature biotechnology, 42(2):275–283.

[17] Holec, P. V., Breuckman, K. C., Leddy, O., White, F. M., Bryson, B. D., and Birnbaum, M. E. (2025). High-throughput screening for class I peptide MHC binding via yeast surface display. Proceedings of the National Academy of Sciences, 122(47):e2514741122.

[18] Hou, C., Liu, D., Zafar, A., and Shen, Y. (2025). Understanding language model scaling on protein fitness prediction. bioRxiv.

[19] Jacobs, T. M., Yumerefendi, H., Kuhlman, B., and Leaver-Fay, A. (2015). SwiftLib: rapid degenerate-codon-library optimization through dynamic programming. Nucleic acids research, 43(5):e34–e34.

[20] Johnston, K. E., Almhjell, P. J., Watkins-Dulaney, E. J., Liu, G., Porter, N. J., Yang, J., and Arnold, F. H. (2024). A combinatorially complete epistatic fitness landscape in an enzyme active site. Proceedings of the National Academy of Sciences, 121(32):e2400439121.

[21] Karp, R. M., Luby, M., and Madras, N. (1989). Monte-carlo approximation algorithms for enumeration problems. Journal of algorithms, 10(3):429–448.

[22] Kingma, D. P. and Ba, J. (2015). Adam: A method for stochastic optimization. In International Conference on Learning Representations (ICLR).

[23] Kloek, T. and van Dijk, H. K. (1978). Bayesian Estimates of Equation System Parameters: An Application of Integration by Monte Carlo. Econometrica, 46(1):1–19.

[24] Krause, A. and Golovin, D. (2014). Submodular function maximization. In Tractability: Practical Approaches to Hard Problems. Cambridge University Press.

[25] Lin, Z., Akin, H., Rao, R., Hie, B., Zhu, Z., Lu, W., Smetanin, N., Verkuil, R., Kabeli, O., Shmueli, Y., et al. (2023). Evolutionary-scale prediction of atomic-level protein structure with a language model. Science, 379(6637):1123–1130.

[26] Marden, J. (2008). Distributed welfare games. Operations Research.

[27] Mason, D., Friedensohn, S., Weber, C., Jordi, C., Wagner, B., Meng, S., Ehling, R., Bonati, L., Dahinden, J., Gainza, P., Correia, B., and Reddy, S. (2021). Optimization of therapeutic antibodies by predicting antigen specificity from antibody sequence via deep learning. Nature Biomedical Engineering, 5:1–13.

[28] Mason, D. M., Weber, C. R., Parola, C., Meng, S. M., Greiff, V., Kelton, W. J., and Reddy, S. T. (2018). High-throughput antibody engineering in mammalian cells by CRISPR/Cas9-mediated homology-directed mutagenesis. bioRxiv.

[29] McInnes, L., Healy, J., and Melville, J. (2018). Umap: Uniform manifold approximation and projection for dimension reduction. arXiv preprint arXiv:1802.03426.

[30] Mena, M. A. and Daugherty, P. S. (2005). Automated design of degenerate codon libraries. Protein Engineering Design and Selection, 18(12):559–561.

[31] O’Donnell, T. J., Rubinsteyn, A., and Laserson, U. (2020). MHCflurry 2.0: Improved pan-allele prediction of MHC class I-presented peptides by incorporating antigen processing. Cell Systems, 11(1):42–48.e7.

[32] Papkou, A., Garcia-Pastor, L., Escudero, J. A., and Wagner, A. (2023). A rugged yet easily navigable fitness landscape. Science, 382(6673).

[33] Passaro, S., Corso, G., Wohlwend, J., Reveiz, M., Thaler, S., Somnath, V. R., Getz, N., Portnoi, T., Roy, J., Stark, H., et al. (2025). Boltz-2: Towards accurate and efficient binding affinity prediction. BioRxiv.

[34] Podgornaia, A. I. and Laub, M. T. (2015). Pervasive degeneracy and epistasis in a protein-protein interface. Science, 347(6222):673–677.

[35] Romero, P. A., Krause, A., and Arnold, F. H. (2013). Navigating the protein fitness landscape with Gaussian processes. Proceedings of the National Academy of Sciences, 110(3):E193–E201.

[36] Salazar, J., Liang, D., Nguyen, T. Q., and Kirchhoff, K. (2020). Masked language model scoring. In Proceedings of the 58th annual meeting of the association for computational linguistics, pages 2699–2712.

[37] Shimko, T. C., Fordyce, P. M., and Orenstein, Y. (2020). DeCoDe: degenerate codon design for complete protein-coding DNA libraries. Bioinformatics, 36(11):3357–3364.

[38] Shuai, R. W., Ruffolo, J. A., and Gray, J. J. (2023). IgLM: Infilling language modeling for antibody sequence design. Cell systems, 14(11):979–989.

[39] Sinai, S., Wang, R., Whatley, A., Slocum, S., Locane, E., and Kelsic, E. (2020). Adalead: A simple and robust adaptive greedy search algorithm for sequence design. arXiv preprint.

[40] Swanson, E., Nichols, M., Ravichandran, S., and Ogden, P. (2025). mBER: Controllable de novo antibody design with million-scale experimental screening. bioRxiv.

[41] Vanchinathan, H. P., Marfurt, A., Robelin, C.-A., Kossmann, D., and Krause, A. (2015). Discovering valuable items from massive data. In Proceedings of the 21th ACM SIGKDD International Conference on Knowledge Discovery and Data Mining, pages 1195–1204.

[42] Vazquez-Lombardi, R., Jung, J. S., Schlatter, F. S., Mei, A., Mantuano, N. R., Bieberich, F., Hong, K.-L., Kucharczyk, J., Kapetanovic, E., Aznauryan, E., Weber, C. R., Zippelius, A., Läubli, H., and Reddy, S. T. (2022). High-throughput T cell receptor engineering by functional screening identifies candidates with enhanced potency and specificity. Immunity, 55(10):1953–1966.e10.

[43] Verma, D., Grigoryan, G., and Bailey-Kellogg, C. (2018). Pareto optimization of combinatorial mutagenesis libraries. IEEE/ACM transactions on computational biology and bioinformatics, 16(4):1143–1153.

[44] Weinstein, E. N., Amin, A. N., Grathwohl, W. S., Kassler, D., Disset, J., and Marks, D. (2022). Optimal design of stochastic DNA synthesis protocols based on generative sequence models. In International Conference on Artificial Intelligence and Statistics, pages 7450–7482. PMLR.

[45] Weinstein, E. N., Gollub, M. G., Slabodkin, A., Gardner, C. L., Dobbs, K., Cui, X.-B., Amin, A. N., Church, G. M., and Wood, E. B. (2024). Manufacturing-aware generative model architectures enable biological sequence design and synthesis at petascale. bioRxiv.

[46] Whitehead, T. A., Chevalier, A., Song, Y., Dreyfus, C., Fleishman, S. J., De Mattos, C., Myers, C. A., Kamisetty, H., Blair, P., Wilson, I. A., et al. (2012). Optimization of affinity, specificity and function of designed influenza inhibitors using deep sequencing. Nature biotechnology, 30(6):543–548.

[47] Williams, R. J. (1992). Simple statistical gradient-following algorithms for connectionist reinforcement learning. Machine learning, 8(3):229–256.

[48] Wu, N. C., Dai, L., Olson, C. A., Lloyd-Smith, J. O., and Sun, R. (2016). Adaptation in protein fitness landscapes is facilitated by indirect paths. elife, 5:e16965.

[49] Wu, N. C., Grande, G., Turner, H. L., Ward, A. B., Xie, J., Lerner, R. A., and Wilson, I. A. (2017). In vitro evolution of an influenza broadly neutralizing antibody is modulated by hemagglutinin receptor specificity. Nature communications, 8(1):15371.

[50] Wu, T. T., Johnson, G., and Kabat, E. A. (1993). Length distribution of CDRH3 in antibodies. Proteins: Structure, Function, and Bioinformatics, 16(1):1–7.

[51] Wu, Z., Kan, S. J., Lewis, R. D., Wittmann, B. J., and Arnold, F. H. (2019). Machine learning-assisted directed protein evolution with combinatorial libraries. Proceedings of the National Academy of Sciences, 116(18):8852–8858.

[52] Yang, J., Ducharme, J., Johnston, K. E., Li, F.-Z., Yue, Y., and Arnold, F. H. (2023). Decoil: Optimization of degenerate codon libraries for machine learning-assisted protein engineering. ACS Synthetic Biology, 12(8):2444–2454.

[53] Yang, K. K., Chen, Y., Lee, A., and Yue, Y. (2019a). Batched stochastic bayesian optimization via combinatorial constraints design. In The 22nd International Conference on Artificial Intelligence and Statistics, pages 3410–3419. PMLR.

[54] Yang, K. K., Wu, Z., and Arnold, F. H. (2019b). Machine-learning-guided directed evolution for protein engineering. Nature methods, 16(8):687–694.

[55] Yoshikawa, A. M., Rangel, A. E., Zheng, L., Wan, L., Hein, L. A., Hariri, A. A., Eisenstein, M., and Soh, H. T. (2023). A massively parallel screening platform for converting aptamers into molecular switches. Nature communications, 14(1):2336.

[56] Zhu, D., Brookes, D. H., Busia, A., Carneiro, A., Fannjiang, C., Popova, G., Shin, D., Donohue, K. C., Lin, L. F., Miller, Z. M., et al. (2024). Optimal trade-off control in machine learning– based library design, with application to adeno-associated virus (AAV) for gene therapy. Science Advances, 10(4):eadj3786.

